# Pneumonic plague protection induced by a monophosphoryl lipid A decorated *Yersinia* outer-membrane-vesicle vaccine

**DOI:** 10.1101/2023.08.17.553697

**Authors:** Saugata Majumder, Shreya Das, Peng Li, Nicole Yang, Hazel Dellario, Haixin Sui, Ziqiang Guan, Wei Sun

## Abstract

A newly constructed *Yersinia pseudotuberculosis* mutant (YptbS46) carrying the *lpxE* insertion and *pmrF-J* deletion exclusively synthesized an adjuvant form of lipid A, monophosphoryl lipid A (MPLA). Outer membrane vesicles (OMVs) isolated from YptbS46 harboring an *lcrV* expression plasmid, pSMV13, were designated OMV_46_-LcrV, which contained MPLA and high amounts of LcrV and displayed low activation of Toll-like receptor 4 (TLR4).

Similar to the previous OMV_44_-LcrV, intramuscular prime-boost immunization with 30 µg of OMV_46_-LcrV exhibited substantially reduced reactogenicity and conferred complete protection to mice against a high-dose of respiratory *Y. pestis* challenge. OMV_46_-LcrV immunization induced robust adaptive responses in both lung mucosal and systemic compartments and orchestrated innate immunity in the lung, which were correlated with rapid bacterial clearance and unremarkable lung damage during *Y. pestis* challenge. Additionally, OMV_46_-LcrV immunization conferred long-term protection. Moreover, immunization with reduced doses of OMV_46_-LcrV exhibited further lower reactogenicity and still provided great protection against pneumonic plague. Our studies strongly demonstrate the feasibility of OMV_46_-LcrV as a new type of plague vaccine candidate.

## Introduction

Plague is a notorious disease in human history caused by *Yersinia pestis* [1]. In the present era, plague is rare, with only a few thousand cases reported annually worldwide, but it may spread, eventually becoming endemic or pandemic due to the presence of large reservoirs in wildlife, expansion of human territory, and turbulence of the current world [2]. Since *Y. pestis* has deadly characteristics, such as high transmissibility and lethality in pulmonary infection and the potential to be used as a biological weapon, it is continuously categorized as a tier 1 select agent by The Centers for Disease Control and Prevention (CDC) [3, 4]. Among the different forms of plague, the primary pneumonic plague caused by direct inhalation of *Y. pestis* rapidly progresses to death within 72 hours if not treated promptly with antibiotics [5]. Moreover, the appearance of multiple drug-resistant *Y. pestis* strains in plague outbreak regions poses severe concerns about the effectiveness of antibiotic treatment [6–8]. Vaccination is the most effective way of preventing infectious diseases, but no fully licensed plague vaccines are available in western countries [4, 9]. Thus, stockpiles of an effective plague vaccine for emergency use are imperative.

Currently, several vaccine candidates based on *Y. pestis* LcrV and/or F1 antigens confer protection in preclinical and clinical studies, including a *Yersinia* outer membrane vesicle (OMV) vaccine [10–15]. In our previous study, a recombinant *Y. pseudotuberculosis* (Yptb) strain, [YptbS44(pSMV13)], carrying the chromosomal insertion of *lpxE* encoding lipid A 1-phosphatase and an *lcrV* expression plasmid was constructed to produce high amounts of self-adjuvating OMVs (designated OMV_44_-LcrV). The self-adjuvating nanoparticles enclosed large amounts of LcrV antigen and exhibited preeminent immunogenicity. However, OMV_44_-LcrV has some obvious drawbacks, such as intramuscular (IM) vaccination resulting in significant animal weight loss, retardation of animal weight gain, high levels of serum IL-1β and IL-6 production, and reactogenicity [10]. Mass spectrometry (MS) analysis revealed that OMV_44_-LcrV still contained a significant proportion of toxic hexa-acylated bisphosphorylated lipid A species [10], which might be a major cause of the above side effects.

The *pmrHFIJKLM* (also termed *arnBCADTEF*) operon in bacteria is responsible for the synthesis and transfer of 4-amino-4-deoxy-L-arabinose (Ara4N), a cationic monosaccharide, to the 1 and 4’-phosphate groups of lipid A [16–18]. In addition, the phosphate groups of lipid A can be glycosylated with Ara4N when *Y. pestis* is grown at a low temperature (less than 30 °C), whereas the Ara4N moiety is scarce or completely absent when *Y. pestis* is cultured at 37 °C [19–21]. In Yptb, the presence of an intact *pmrHFIJKLM* operon [22] contributes to Ara4N addition to the phosphate groups in lipid A of Yptb and its OMVs [10]. The *lpxE* expression in the YptbS44(pSMV13) strain only partially removed the 1-phosphate group from lipid A species [10]. Therefore, we speculate that the presence of Ara4N may hinder the dephosphorylation of lipid A by LpxE. Disruption of Ara4N synthesis in YptbS44(pSMV13) may promote the production of monophosphoryl lipid A species, which would reduce reactogenicity but retain the immunogenicity of OMVs from the corresponding mutant. In this study, a new mutant strain, YptbS46, was constructed by introducing a *pmrF-J* mutation into the YptbS44 strain and shown to exclusively produce monophosphoryl lipid A (MPLA) species. Then, OMV_46_-LcrV isolated from the YptbS46 harboring the pSMV13 plasmid demonstrated significantly low reactogenicity and retained excellent immunogenicity in protection against pneumonic plague. OMV_46_-LcrV vaccination induced robust humoral and cellular immune responses, orchestrated innate immunity in the lung, and conferred short- and long-term protection against high doses of pulmonary *Y. pestis* infection.

## Results

### Construction of a recombinant *Y. pseudotuberculosis* strain exclusively producing monophosphoryl lipid A-decorated OMVs

We assumed that a *pmrF-J* mutation introduced into the YptbS44 strain to generate a new strain, YptbS46 (Table S1), could disrupt Ara4N synthesis and promote comprehensive production of monophosphoryl lipid A (MPLA) species. Mass spectrometry analysis revealed that the YptbS46 strain carrying Δ*pmrF-J* failed to synthesize the Ara4N moiety and exclusively produced monophosphoryl lipid A species in comparison to the YptbS44 strain (Fig. 1A). Then, plasmids pSMV13 and pYA3493 (an empty plasmid) (Table S1) were introduced into the YptbS46 mutant to determine LcrV synthesis in bacteria and their OMVs, respectively (Fig. 1B). High amounts of LcrV antigen were synthesized in the YptbS46(pSMV13) strain and enclosed in its OMVs but not in the YptbS46(pYA3493) strain and its OMVs. To simplify the term OMVs in this context, OMVs from YptbS46(pSMV13) and YptbS46(pYA3493) were designated OMV_46_-LcrV and OMV_46_-NA, respectively.

**Figure. 1.**
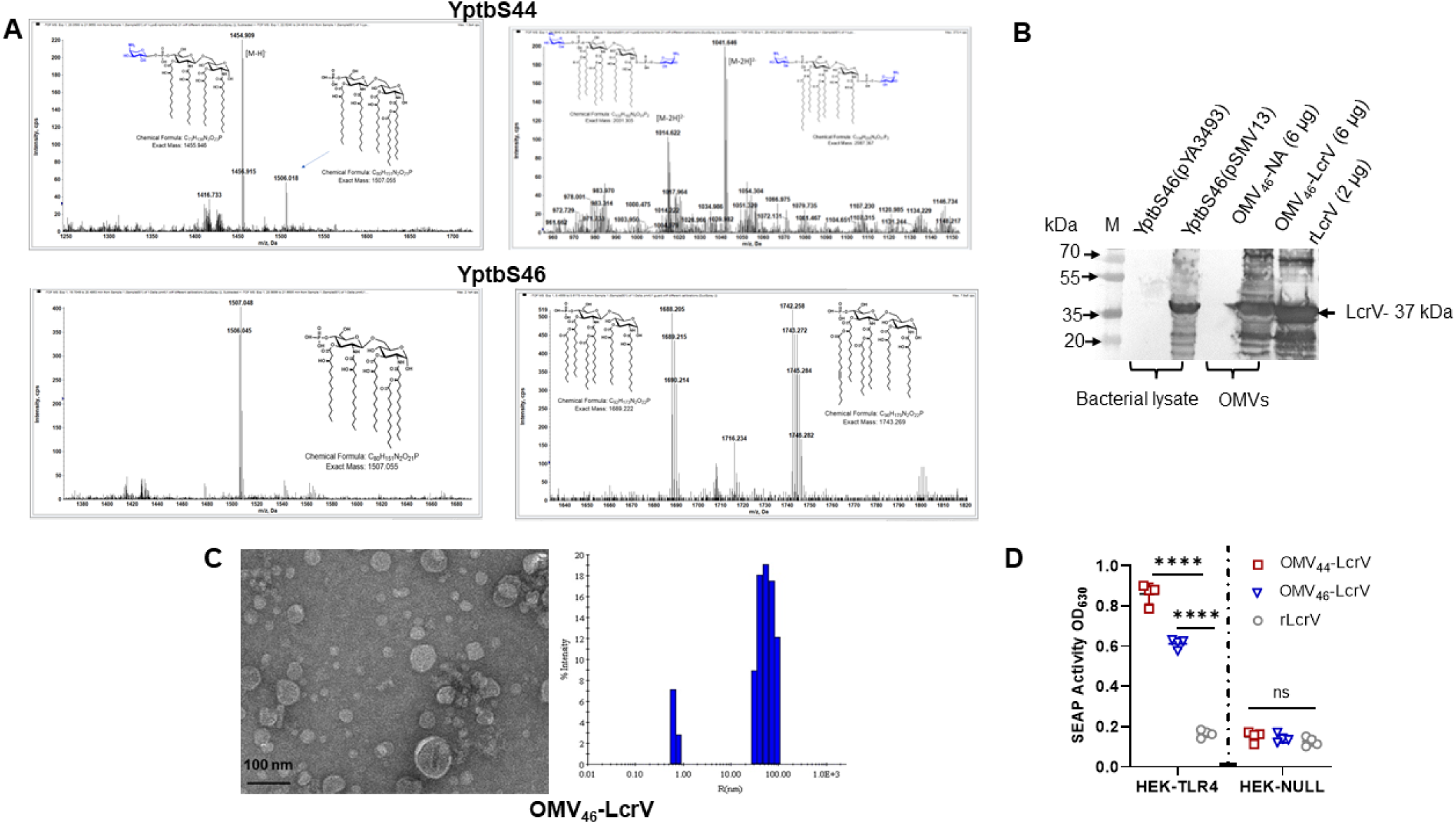
Lipid A profiles of recombinant Yptb strains and analysis of OMVs from these strains. (A) Mass spectrometry analysis of lipid A species in the YptbS44 and YptbS46 strains. Shown are the negative ion electropray ionization (ESI) mass spectra of the [M-H]^-^ ions of lipid A species and their chemical structures. (B) Analysis of LcrV antigen by immunoblotting. YptbS46(pYA3493): YptbS46 harboring an empty Asd^+^ plasmid; YptbS46(pSMV13): YptbS46 harboring an Asd^+^ plasmid for *lcrV* expression; OMV_46_-NA: 6 µg of OMVs isolated from YptbS46(pYA3493); OMV_46_-LcrV: 6 µg of OMVs isolated from YptbS46(pSMV13); 2 µg of purified rLcrV used as a control; M: 10-250 kDa protein marker (Thermo Fischer Scientific). (C) Transmission electron microscopy (TEM) image of OMV-LcrV (left panel) and dynamic light scattering (DLS) of OMV-LcrV (right panel). (D) Comparison of embryonic alkaline phosphate (SEAP) activities in HEK-blue cells with or without mTLR4. HEK-blue mTLR4 cells were cocultured with 40 µg/mL OMVs for 8 hours. OMV_44_-LcrV from YptbS44(pSMV13) and rLcrV were used as controls. Data are shown as the mean ± SD. The statistical significance of differences among groups was analyzed by two-way ANOVA with Tukey’s post hoc test: ns, no significance; **** *p<0.0001*.

Next, the morphology and size of OMV_46_-LcrV were characterized by transmission electron microscopy (TEM) and dynamic light scattering (DLS). OMVs showed a circular morphology with a bilayer structure and were in the range from 20 to 200 nm (Fig. 1C). Furthermore, we compared the secreted embryonic alkaline phosphatase (SEAP) activity of HEK-blue mTLR4 cells cultured with different OMVs (Fig. 1D). The results showed that the TLR4 stimulatory activity of OMV_46_-LcrV was significantly less than that of OMV_44_-LcrV constructed previously [10], whereas it was substantially higher than that of purified LcrV protein. No SEAP activity was detected in HEK-blue null cells with the same stimulation. Thus, the exclusive monophosphoryl lipid A modification diminished the TLR4 activation of OMV_46_-LcrV.

### Protection induced by OMV immunization against pneumonic plague

Groups of Swiss Webster mice (n=10/group, equal males and females) were intramuscularly immunized with 30 µg of different OMVs (OMV_44_-LcrV, OMV_46_-LcrV, and OMV_46_-NA), 3 µg of rLcrV/Alhydrogel, or 1x sterile phosphate-buffered saline (PBS, sham) as a negative control and then boosted on day 22 after priming immunization (Fig. 2A). Similar to previous observations [10], OMV_44_-LcrV immunization resulted in ~9% weight loss at 4 days post-vaccination (DPV) and substantially retarded animal weight gain during the entire immunization (Fig. 2B), as well as swelling at the site of injection and anorexia until 3 DPV (observation data). However, mice vaccinated with OMV_46_-LcrV or -NA displayed significantly less weight loss (4-5%) on 4 DPV and then rapidly gained weight (Fig. 2B). No significant swelling at the site of injection or anorexia was observed in the OMV_46-_immunized mice (observation data). Furthermore, the amounts of serum IL-6 and IL-1β elicited in OMV_46_-LcrV-immunized mice were significantly less than those in OMV_44_-LcrV-immunized mice but still substantially higher than those in rLcrV-immunized or sham mice at 1 and 3 DPV (Fig. 2C).

**Figure 2:**
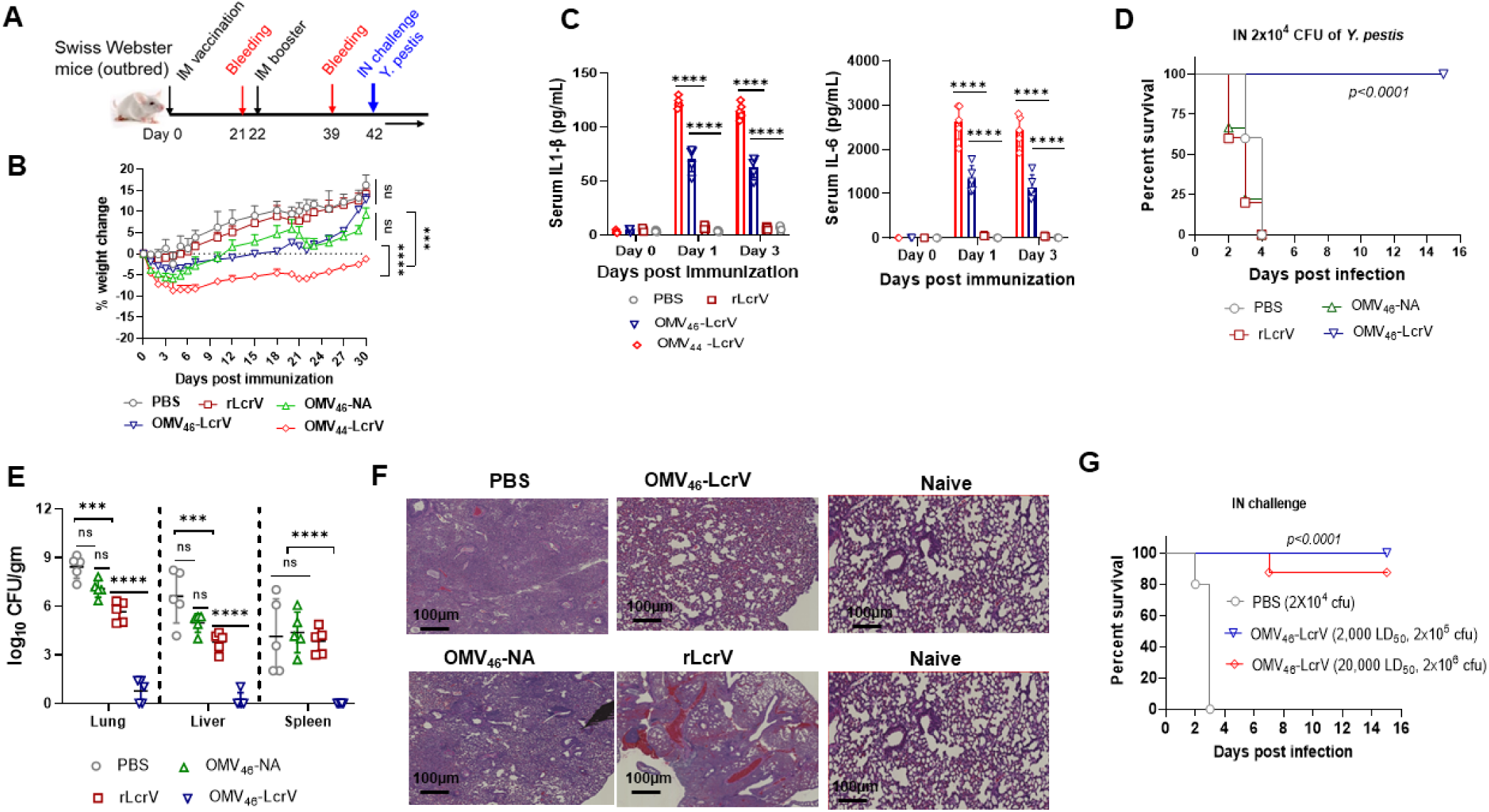
Protective efficacy of intramuscular immunization (IM) with OMVs against pneumonic plague. Swiss Webster mice (n=10, equal males and females, 7 weeks old) were immunized IM with 30 µg of OMV_44_-LcrV, OMV_46_-LcrV, OMV_46_-NA, 3 µg of rLcrV/Alhydrogel, or Alhydrogel alone in 50 µL of PBS and then boosted on day 22 after priming. (A) Immunization schema for the animal study. (B) Weight change rates of animals post-immunization. (C) Analysis of serum cytokine levels in immunized mice. Cytokines (IL-1β and IL-6) in the sera collected on 0, 1, and 3 DPV were analyzed using corresponding ELISA kits (Invitrogen). (D) On 42 DPV, mice were intranasally challenged with 2×10^4^ CFU (200 LD_50_) of *Y. pestis* KIM6+(pCD1Ap). (E) At 2 DPI, the bacterial burden was determined in the lung, liver, and spleen. The experiments were performed twice, and the data were combined for analysis. (F) Lung histopathological analysis of representative mice from each group at 2 DPI. The lungs were microscopically examined and imaged using a Nanozoomer 2.0 RS Hamamatsu slide scanner (scale bar, 100 nm). (G) Swiss Webster mice immunized with OMV_46_-LcrV were intranasally challenged with 2×10^5^ CFU (2,000 LD_50_) and 2×10^6^ CFU (20,000 LD_50_) of *Y. pestis* KIM6+(pCD1Ap). After infection, animal morbidity and mortality were monitored for 15 days. Statistical significance was analyzed by the log-rank (Mantel-Cox) test for survival analysis. Data were analyzed and presented as the mean ± SD. The statistical significance of differences among groups was analyzed by two-way ANOVA with Tukey’s post hoc test: ns, no significance; *, *P*< *0.05*; **, *P*< *0.01*; ****, *P*< *0.0001*.

Similar to the OMV_44_-LcrV vaccination [10], the OMV_46_-LcrV vaccination afforded complete protection against an intranasal (IN) challenge with 2×10^4^ CFU (200 LD_50_) of *Y. pestis* KIM6+(pCD1Ap), while OMV_46-_NA, LcrV, or sham immunization failed to provide any protection against the same challenge (Fig 2D). At 2 days post-infection (DPI), the lungs of sham mice had extremely high *Y. pestis* titers (mean 8.4 log_10_ CFU/g tissue). *Y. pestis* disseminated into the liver (mean 6.6 log_10_ CFU/g tissue) and spleen (mean 4.1 log_10_ CFU/g tissue) at significant bacterial titers (Fig. 2E). In LcrV- or OMV_46_-NA-vaccinated mice, the lungs and liver had moderate and reduced bacterial titers, while the spleen had comparable bacterial titers to those of sham mice (Fig. 2E). Very low titers of *Y. pestis* were detected in the lungs, liver, and spleen of OMV_46_-LcrV-vaccinated mice (Fig. 2E). Histopathological analysis showed that mild and transient immune cell infiltration and inflammation were observed in the lungs of OMV_46_-LcrV-immunized mice at 2 DPI in comparison to the naïve lungs. However, large amounts of immune cell infiltration, hemorrhage, edema, and congestion were observed in the lungs of sham-, LcrV-, or OMV_46_-NA-immunized mice (Fig. 2F). Moreover, the OMV_46_-LcrV-immunized mice intranasally challenged with 2,000 LD_50_ and 20,000 LD_50_ of *Y. pestis* had 100% and 80% survival, respectively (Fig. 2G).

To further reduce reactogenicity, we also generated YptbS47 (Table S1) on top of YptbS46 by deletion of *lpxL* encoding a lauroyltransferase that involves the addition of the secondary lauroyl residue to the distal glucosamine unit [23]. Mass spectrometry analysis showed that the YptbS47 strain lacked the secondary lauroyl residue of lipid A species (Fig. S1A). OMV_47_-LcrV from YptbS47(pSMV13) exhibited even lower TLR4 stimulation *in vitro* and reactogenicity *in vivo* than OMV_46_-LcrV (Fig. S1B and C). The OMV_47_-LcrV-immunized mice intranasally challenged with 200 LD_50_ and 2,000 LD_50_ of *Y. pestis* had complete and 60% survival, respectively (Fig. S1D). Given the above data, we chose OMV_46_-LcrV manifesting superimmunogenicity and relatively low reactogenicity for further studies.

### Antibody and T-cell responses induced by OMVs

High serum anti-LcrV IgG titers were elicited in both LcrV- and OMV_46_-LcrV-immunized mice at 21 DPV and boosted at 39 DPV. However, anti-LcrV IgG titers in OMV_46_-LcrV-immunized mice were substantially higher than those in LcrV-immunized mice at both time points (Fig. 3A). No anti-LcrV IgG titers were detected in sham or OMV_46_-NA-immunized mice (Fig. 3A). Antibody isotyping demonstrated that both ratios of anti-LcrV IgG2a/IgG1 and IgG2b/IgG1 in OMV_46-_LcrV-immunized mice were almost equal to one at both 21 and 39 DPV and were significantly higher than the ratios (mean value ~0.8) in LcrV-immunized mice (Fig. 3B). Thus, OMV_46-_LcrV immunization induced a balanced Th1/Th2 response, while LcrV immunization induced a Th2-biased response. In addition, significant amounts of anti-LcrV IgG titers in the bronchoalveolar lavage fluid (BALF) were induced in OMV_46-_LcrV-immunized mice but not in rLcrV-immunized mice (Fig. S2A). No LcrV-specific secretory IgA (sIgA) was detected in the BALF of OMV_46-_LcrV- or rLcrV-immunized mice (data not shown).

**Figure 3:**
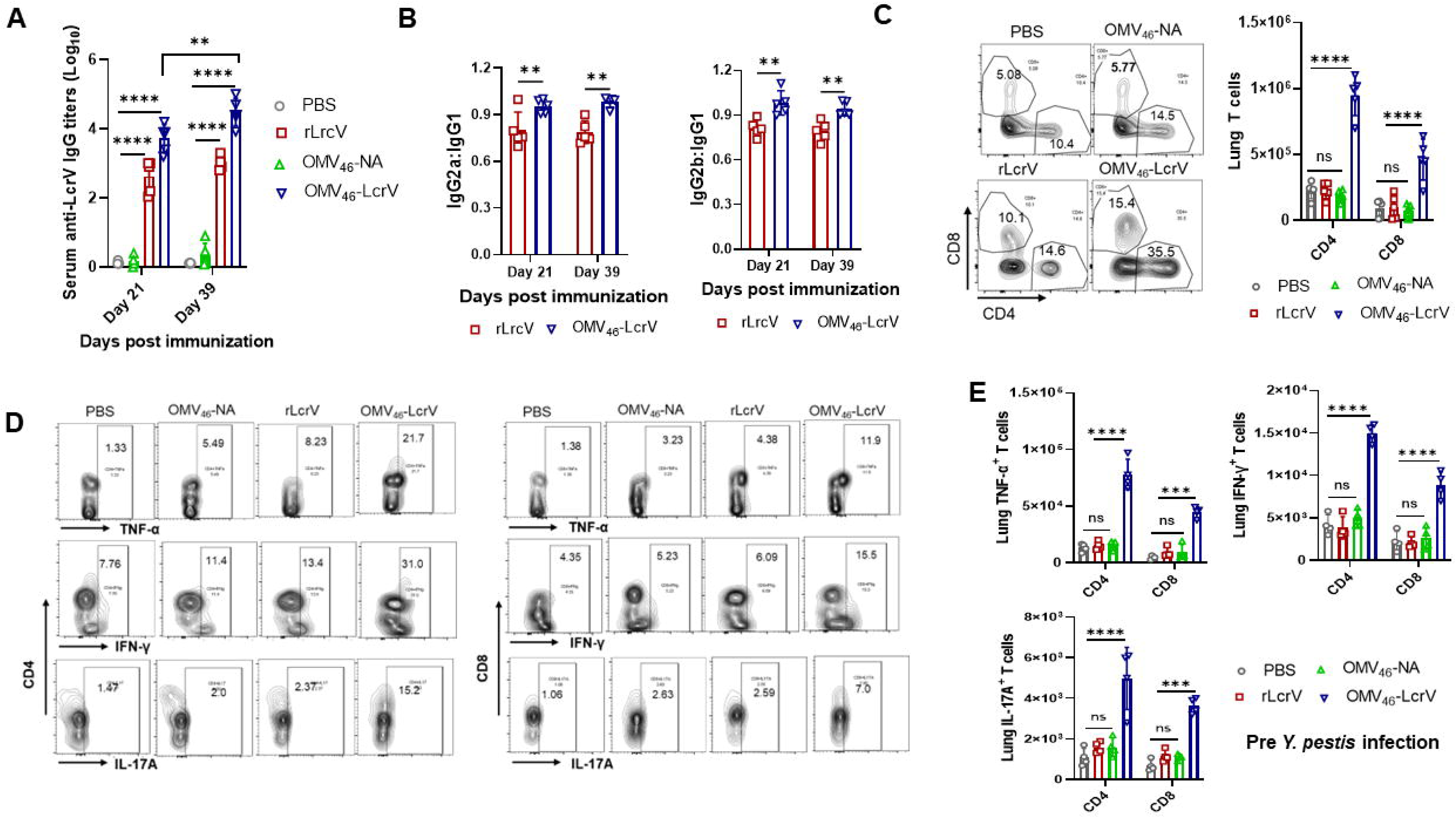
Antibody and T-cell responses in the immunized mice. Swiss Webster mice (n=5, equal males and females, 7 weeks old) were immunized IM with 30 µg of OMV_46_-LcrV, OMV_46_-NA, 3 µg of rLcrV/Alhydrogel or Alhydrogel alone in 50 µL of PBS and boosted on day 22 after priming. (A) Total serum IgG titers to LcrV in Swiss Webster mice at 21 and 39 DPV. (B) The ratio of IgG2a/IgG1 and IgG2b/IgG1 for antibodies specific to LcrV antigen on 21 and 39 DPV. (C) On 42 DPV, lymphocytes were aseptically isolated from the lungs and stimulated *in vitro* with 20 μg/mL rLcrV protein for 48 h to identify antigen-specific CD4^+^ and CD8^+^ T cells. Lung cells from sham mice were used as a control. Flow plots (left) and quantitative analysis (right) of total lung CD4^+^ and CD8^+^ T cells in the postimmunization. (D) Flow plots and (E) quantitative analysis of total CD4^+^ and CD8^+^ T cells producing IFN-γ, TNF-α, or IL-17A cytokines from the lungs. Each symbol was obtained from an individual mouse, and data are represented as the mean ± SD. The statistical significance of differences among groups was analyzed by two-way ANOVA with Tukey’s post hoc test: ns, no significance; * *p<0.05*; ** *p<0.01*; *** *p<0.001*; **** *p<0.0001*.

To determine antigen-specific T-cell responses, lung single cells were isolated from immunized mice at 42 DPV and stimulated *in vitro* with rLcrV for 48 h, stained with the indicated antibodies, and analyzed using flow cytometry. Fig. S2B shows the gating strategy. Flow and quantitative plots showed that the number of both LcrV-specific lung CD4^+^ and CD8^+^ T cells from OMV_46_-LcrV-immunized mice was significantly higher than that of sham-, LcrV-, or OMV-NA-immunized mice (Fig. 3C). Similar profiles were observed in the number of lung CD4^+^ and CD8^+^ T cells producing interferon-gamma (IFN-γ), tumor necrosis factor (TNF-α), or interleukin 17A (IL-17A) (Figs. 3D and E). In addition, similar patterns of LcrV-specific T cells were observed in the spleen (Fig. S2C).

Moreover, lung T-cell responses *in vivo* were evaluated in immunized mice upon pulmonary *Y. pestis* infection. On 42 DPV, mice were intranasally challenged with 200 LD_50_ of *Y. pestis*. At 2 DPI, the lung and spleen were isolated from the immunized and PBS mice to determine the total T-cell number and specific cytokines by flow cytometry. Similar to *ex vivo* studies (Figs. 3C-E), the number of CD4^+^ and CD8^+^ T cells and corresponding cells producing cytokines (IFN-γ, IL-17A, and TNF-α) in the lung (Fig. 4A) and spleen (Fig. S2D) of OMV_46_-LcrV-immunized mice was substantially higher than that in sham-, LcrV-, or OMV_46_-NA-immunized mice. Taken together, robust antibody and T-cell responses in both lung mucosal and systemic compartments elicited by OMV_46-_LcrV immunization may contribute to remarkable protection against pneumonic plague.

**Figure 4:**
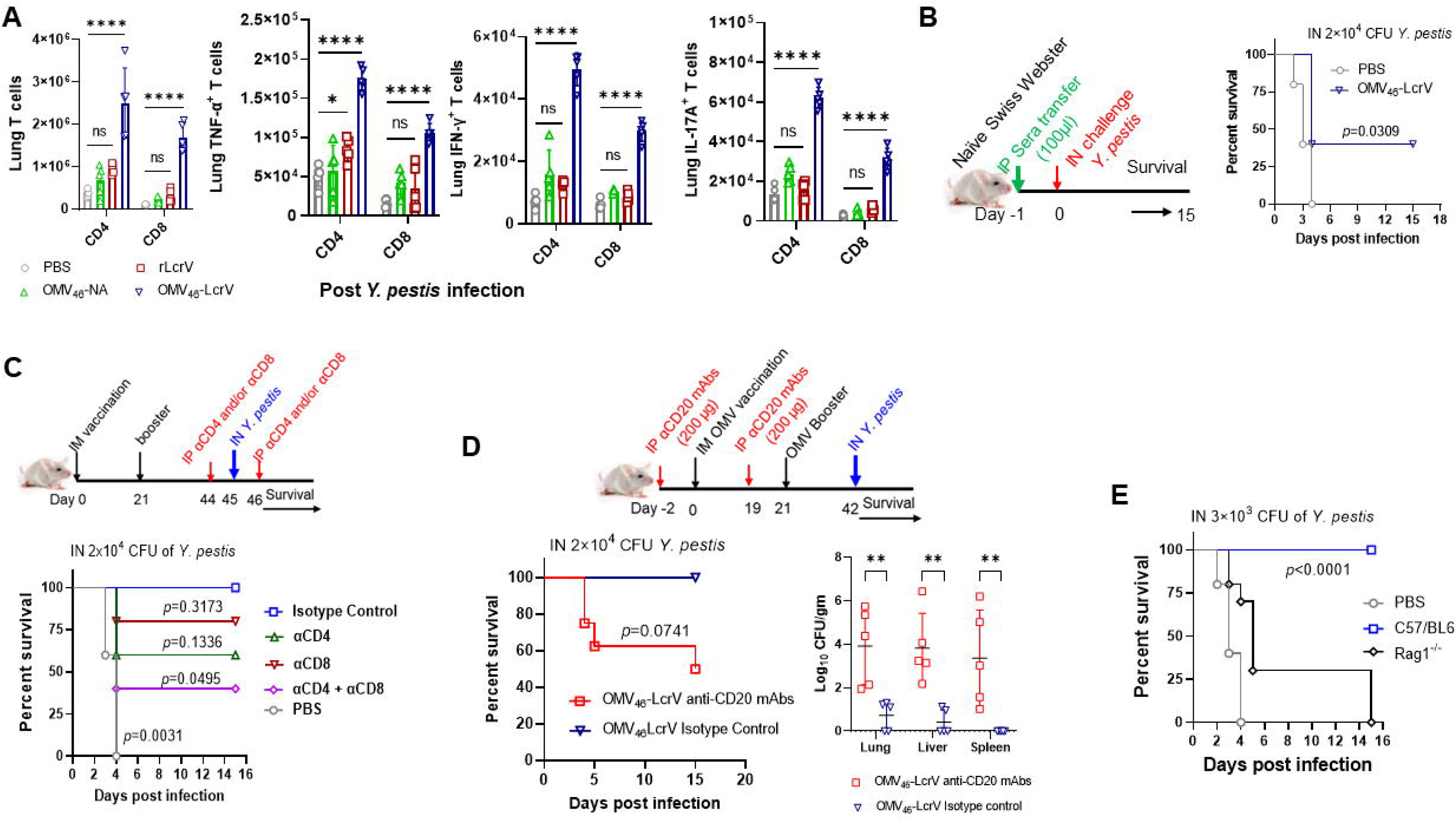
The role of adaptive immune responses in protection against pneumonic plague. (A) Quantitative analysis of lung CD4^+^ and CD8^+^ T cells and their corresponding IFN-γ-, TNF-α-, or IL-17A-producing cells in immunized mice after pulmonary *Y. pestis* infection. On 42 DPV, mice were intranasally challenged with 2×10^4^ CFU (200 LD_50_) of *Y. pestis* KIM6+(pCD1Ap). At 2 DPI, lung cells were harvested from euthanized Swiss Webster mice (n=5/group females) and stained for T cells and specific cytokines (IFN-γ, TNF-α, or IL-17A) for flow cytometry. (B) Protection of serum transfer. (Left) The schema of serum transfer and (Right) naïve Swiss Webster mice (n=5/group females) were IP injected with 100 µl of sera from sham and OMV_46_-LcrV-immunized mice collected on 35 DPV. Twenty-four hours postinjection, recipient mice were intranasally challenged with 200 LD_50_ of *Y. pestis* KIM6+(pCD1Ap). (C) Protection of T-cell depletion. (Up) The schema of T-cell depletion and OMV_46_-LcrV-immunized Swiss Webster mice (n=5/group females) were IP administered either with anti-CD4, anti-CD8, or anti-CD4 plus anti-CD8 mAbs (200 µg/each mouse in 200 µl PBS) at a two-day interval for the depletion of CD4^+^ and/or CD8^+^ T cells, mice were injected with the isotype control mAbs as controls. The treated mice were IN challenged with 200 LD_50_ of *Y. pestis* KIM6+(pCD1Ap). (Down) Animal survival was monitored for 15 days. (D) Protection of B-cell depletion. (Top) Schema for B-cell depletion before OMV_46_-LcrV immunization. Swiss Webster mice (n=5-7/group, females) were IP administered anti-CD20 mAbs on day 2 before immunization and again treated with the same antibody on day 19 (two days before the booster at day 21) for the depletion of B cells. Mice were injected with isotype control mAbs as controls. Treated mice were IN challenged with 2.0×10^4^ CFU of *Y. pestis* KIM6+(pCD1Ap). (Lower left) Animal survival was monitored for 15 days. (Lower right) Bacterial burden was determined in the lung, liver, and spleen for the B-cell-depleted and isotype antibody-treated control mice at 2 DPI. The experiments were performed twice, and the data were combined for analysis. (E) Protection in B and T-cell-deficient mice against pneumonic plague. C57/BL6 and Rag1^−/−^ mice (n=5, mixed gender) were intramuscularly immunized with 20 µg of OMV_46_-LcrV. Mice administered PBS were used as a negative control group. On 42 DPV, animals were intranasally challenged with 30 LD_50_ of *Y. pestis*. The mortality and morbidity of animals were monitored for 15 days. Statistical significance was analyzed by the log-rank (Mantel-Cox) test for survival analysis. Data were analyzed and presented as the mean ± SD. The statistical significance of differences among groups was analyzed by two-way ANOVA with Tukey’s post hoc test: ns, no significance; *, *P*< *0.05*; **, *P*< *0.01*; ****, *P*< *0.0001*.

### Both humoral and cellular responses are required for full protection against pneumonic plague

Given that OMV_46-_LcrV immunization induced robust LcrV-specific antibody and T-cell responses (Figs. 3A-B, 4A, and S2), we sought to determine the roles of antibodies and T cells in protection against pneumonic plague. To ascertain the role of antibodies, immune sera collected from mice were adoptively transferred to naïve mice. Intraperitoneal (IP) injection with 100 µl of sera collected at 39 DPV from OMV_46-_LcrV-immunized mice provided 40% protection, while sera from sham mice failed to offer any protection (Fig. 4B). To ascertain the importance of T cells in protection, groups of OMV_46-_LcrV-immunized mice were intraperitoneally injected with 200 µg of mouse anti-CD4, anti-CD8 monoclonal antibodies (mAbs), or both to deplete specific T cells and then intranasally challenged with *Y. pestis,* as shown in the schema (Figs. 4C up and S3B). Depletion of CD4^+^, CD8^+^, and both T cells reduced animal survival to 60%, 80%, and 40%, respectively, whereas mice injected with isotype control antibodies completely survived the same infection (Figs. S3A and 4C down).

Studies have reported that B cells contribute to T-cell activation and expansion *in vivo* and pathogen control [24, 25]. To further determine the role of the humoral response in protection, groups of Swiss Webster mice were intraperitoneally injected with 200 µg of mouse anti-CD20 mAbs to specifically deplete B cells [24] at 2 days before the OMV_46_-LcrV prime and booster immunization shown in the schema (Fig. 4D top). B-cell depletion was confirmed by flow cytometry (Figs. S3B and C) and showed a significant reduction in serum anti-LcrV IgG titers in OMV_46_-LcrV-immunized mice compared to isotype control-injected mice (Fig. S3D). Then, mice were subjected to a pulmonary challenge with 200 LD_50_ of *Y. pestis*. The results showed that B-cell depletion reduced survival to 50%, whereas the isotype control retained 100% survival (Fig. 4D, bottom left). At 2 DPI, anti-CD20-treated mice also had higher *Y. pestis* CFUs in the lung, spleen, and liver compared to the mice treated with isotype control antibodies (Fig. 4D, bottom right).

Furthermore, C57/BL6 and Rag1^−/−^ mice (n=5-10, mixed males and females) were immunized with OMV_46-_LcrV and then challenged intranasally with 30 LD_50_ of *Y. pestis*. Immunized C57/BL6 mice completely survived the challenge, whereas seven out of ten Rag1^−/−^ mice died within 4 days, and the remaining 3 mice were considered dead because they lost more than 20% weight at 15 DPI. Sham mice succumbed within 3 DPI (Fig. 4E). Collectively, both humoral and cellular responses are required for full protection against pulmonary *Y. pestis* infection.

### Alterations in lung alveolar macrophages and neutrophils in immunized mice pre- and post-pulmonary *Y. pestis* infections

Alveolar macrophages (AMs) are targeted in early pulmonary *Y. pestis* infection, followed by preferential targeting of neutrophils in the later stage [26]. Additionally, the early apoptosis of macrophages modulated by *Y. pestis* was shown to promote the progression of primary pneumonic plague [27]. A hallmark of primary pneumonic plague is massive neutrophil recruitment to the lung during the proinflammatory phase of the disease [28]. Therefore, we sought to examine the number of AMs and neutrophils in the lungs of immunized mice pre- and post-infections as per the gating strategy shown in Fig. S3E. Based on the flow (Fig. 5A) and quantitative plots (Fig. 5B), both OMV_46-_LcrV and OMV_46-_NA immunization significantly increased the number of AMs in both lungs and BALFs compared to sham or LcrV immunization. The number of AMs in the lungs and BALFs was comparable between the sham and LcrV immunization groups (Fig. 5B). Compared with the pre-infection, the pulmonary *Y. pestis* infection significantly increased the number of AMs in the lungs and BALFs of OMV_46-_LcrV-immunized mice but retained comparable numbers in those of the remaining groups of mice (Figs. 5C and D). Simultaneously, lung neutrophil analysis indicated that the number of neutrophils in the lungs and BALFs of all groups of mice was comparable at preinfection (Figs. 5E and F). Compared with preinfection, pulmonary *Y. pestis* infection dramatically increased the number of neutrophils in the lungs and BALFs of sham-, LcrV-, or OMV_46-_NA-immunized mice but retained a comparable number in those of OMV_46-_LcrV-immunized mice (Figs. 5G and H). The results revealed that OMV_46-_LcrV immunization effectively increased activated AMs and contained massive neutrophil influx after pulmonary *Y. pestis* infection, which facilitated bacterial clearance and limited lung damage.

**Figure 5:**
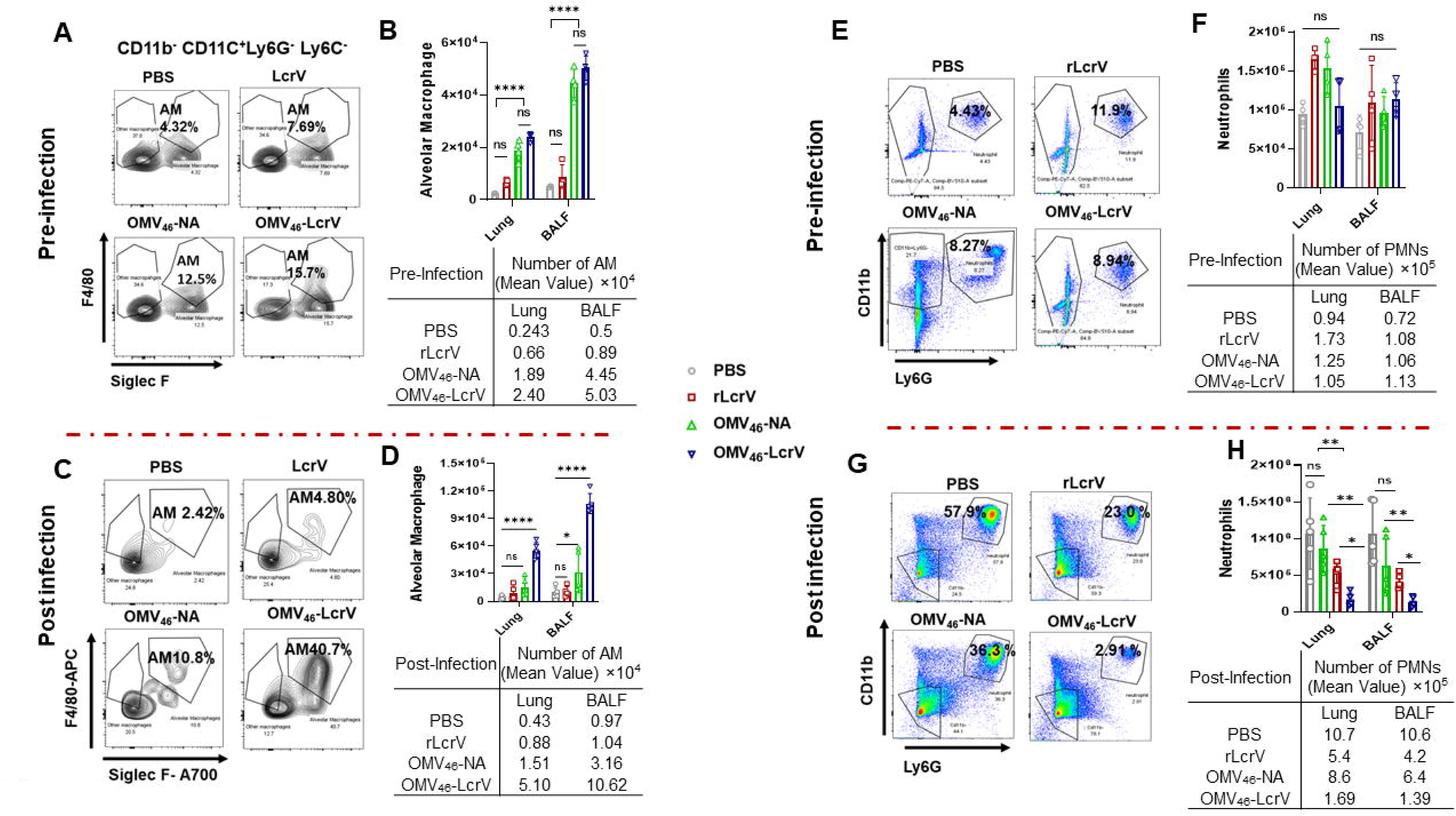
Pattern of alveolar macrophages (AMs) and neutrophils in the lung and BALF with or without infection. Swiss Webster mice (n=5 females) were immunized with OMV_46_-LcrV, OMV_46_-NA, rLcrV, or PBS as described above. Single cells were collected from the lung and BALF at 42 DPV and 2 DPI and stained with fluorescent markers to characterize AMs and neutrophils using flow cytometry. (A) Representative flow plots showing the percentage of AMs and (B) quantitative analysis of the number of AMs in the lung (per lung) and BALF (per/mL) at 42 DPV. (C) Representative flow plot showing the percentage of AMs in the lung (per lung) and BALF (per/mL) at 2 DPI. (D) Quantitative analysis of the number of AMs in the lung (per lung) and BALF (per/mL) at 2 DPI. (E) Representative flow plot showing the percentage of neutrophils in the lung (per lung) and BALF (per/mL) at 42 DPV. (F) Quantitative analysis of the number of neutrophils in the lung (per lung) and BALF (per/mL) at 42 DPV. (G) Representative flow plot showing the percentage of neutrophils in the lung (per lung) and BALF (per/mL) at 2 DPI. (H) Quantitative analysis of the number of neutrophils in the lung (per lung) and BALF (per/mL) at 2 DPI. Data were analyzed and presented as the mean ± SD. The statistical significance of differences among groups was analyzed by two-way ANOVA with Tukey’s post hoc test: ns, no significance; *, *P*< *0.05*; **, *P*< *0.01*; ****, *P*< *0.0001*.

### Evaluation of pneumonic plague protection in long-term and dose-reduced immunized mice

To evaluate the long-term protection of OMV_46-_LcrV immunization, a prime-boost immunization was conducted following the same regimen shown in Fig. 2A, and blood was collected from mice at 60, 90, 120, 150, 180, and 240 DPV to monitor variations in antibody titers (Fig. 6A). High serum anti-LcrV IgG titers retained comparable levels up to 180 DPV without a significant decline but started significantly declining at 210 DPV (Fig. 6B). The anti-LcrV IgG titers at 240 DPV were significantly lower than those from 60 to 150 DPV but still maintained moderate IgG titers (Fig. 6B). On 245 DPV, mice were subjected to an IN challenge with 400 LD_50_ of *Y. pestis*. Eighty percent of OMV_46-_LcrV-immunized mice survived, while no sham mice survived the same challenge (Fig. 6C). At 2 DPI, substantially high titers of *Y. pestis* were detected in the lungs (mean 7.13 log_10_ CFU/g tissue), liver (mean 7.98 log_10_ CFU/g tissue), and spleen (mean 4.83 log_10_ CFU/g tissue) of sham mice (Fig. 6D). Moderate bacterial titers were detected in the lung (mean 2.61 log_10_ CFU/g tissue) and liver (mean 3.46 log_10_ CFU/g tissue), and limited bacteria were detected in the spleen (mean 0.31 log_10_ CFU/g tissue) of OMV_46-_LcrV-immunized mice (Fig. 6D).

**Figure 6:**
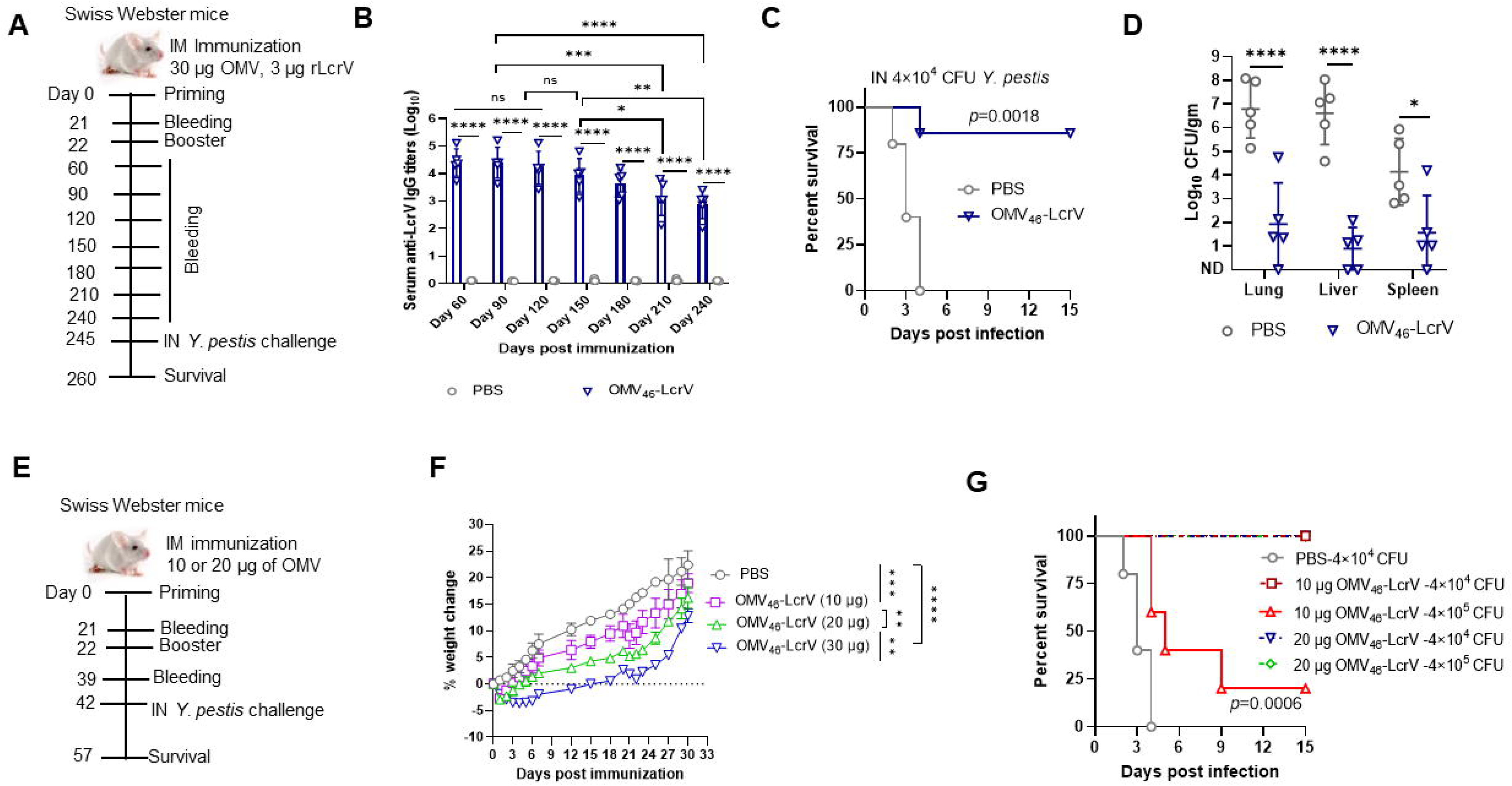
Pneumonic plague protection in long-term and dose-reduced immunized mice. Swiss Webster mice (n=10/group, equal males and females) were immunized intramuscularly with OMV_46_-LcrV or Alhydrogel alone in 50 µL of PBS (negative control) and then boosted on day 22 after the priming immunization as mentioned previously. (A) Schema for long-term protection in mice immunized with 30 µg of OMV_46_-LcrV. (B) Total anti-LcrV IgG titers in sera. Blood was collected from the immunized mice at 60, 90, 120, 150, 180, 210, and 240 DPV. (C) On 245 DPV, the immunized mice were intranasally challenged with 4.0×10^4^ CFU (400 LD_50_) of *Y. pestis* KIM6+(pCD1Ap), and survival was monitored for 15 days. (D) At 2 DPI (i.e., 247 DPV), the bacterial burden was determined in the lung, liver, and spleen. (E) Schema for evaluation of dose-reduction immunization with 20 or 10 µg of OMV_46_-LcrV. (F) Weight change rates of immunized mice. (G) At 42 DPV, the immunized mice were challenged with 400 LD_50_ and 4,000 LD_50_ of *Y. pestis* KIM6+(pCD1Ap). Survival was monitored for 15 days. Statistical significance was analyzed by the log-rank (Mantel-Cox) test for survival analysis. Data were analyzed and presented as the mean ± SD. The statistical significance of differences among groups was analyzed by two-way ANOVA with Tukey’s post hoc test: ns, no significance; *, *P*< *0.05*; **, *P*< *0.01*; ****, *P*< *0.0001*.

The reduced dose of vaccines may have low reactogenicity and increase vaccine coverage in situations of limited global vaccine supply if they can achieve comparable protection as their high-dose counterparts [29]. Given this, we sought to evaluate the protective efficacy of OMV_46_-LcrV using reduced doses. Mice were immunized with 10 or 20 μg of OMV_46-_LcrV following the same regimen as described above. The results showed that the weight loss and gain of mice vaccinated with 10 or 20 μg of OMV_46-_LcrV were significantly better than those of mice vaccinated with 30 μg of OMV_46-_LcrV (Fig. 6F). Moreover, 20 μg of OMV_46-_LcrV immunization afforded complete protection against both 400 and 4,000 LD_50_ of *Y. pestis* pulmonary challenge. Ten μg of OMV_46-_LcrV immunization also provided complete protection against 400 LD_50_ of *Y. pestis* challenge but bare protection against a high-dose challenge (Fig. 6G). Thus, the results demonstrated that OMV_46-_LcrV immunization afforded long-term protection against pneumonic plague, and the immunization dosage was associated with protection in the context of a specific challenge dose.

## Discussion

Bacterial OMVs are complex bionanoparticles that contain immune stimulators [e.g., lipopolysaccharide (LPS), proteins, DNA, etc.] and antigenic molecules that can trigger both innate and adaptive immune responses [30]. The mounted responses are a double-edged sword: they can be either beneficial for counteracting pathogen infections or detrimental to hosts if immune dysregulation is induced by toxic components in the OMVs [31, 32]. Among them, LPS is one of the major components on the surface of OMVs from gram-negative bacteria and causes toxicity to mammalian hosts [31, 33, 34]. Thus, minimizing LPS toxicity in OMVs to prepare a safe and effective OMV vaccine within the tolerance level of humans has been studied for decades [35]. Reduction of LPS toxicity can be achieved by inactivating lipid A biosynthesis acyltransferases, such as LpxL or LpxM, to generate penta- or tetra-acylated lipid A species [36] or by expressing *lpxE* to remove a phosphate group of lipid A [37, 38]. The heterologous expression of *lpxE* in a recombinant *Y. pseudotuberculosis* (YptbS44) strain only partially removed the 1-phosphate group from toxified lipid A (hexa-acylated lipid A) [10]. OMVs from YptbS44 had significantly reduced TLR4 stimulation but still exhibited obvious toxicity and reactogenicity [10]. Here, we demonstrated that loss of L-Ara4N moieties in lipid A via disruption of *pmrF-J* in the YptbS44 strain resulted in exclusive synthesis of monophosphoryl lipid A species, which further reduced TLR4 stimulation and reactogenicity of OMV_46_ from the YptbS46 strain (Figs. 1D, 2A and B). Adding the *lpxL* mutation into the YptbS46 strain to remove one fatty acid chain from MPL further decreased TLR4 stimulation and reactogenicity of OMV_47_-LcrV (Fig. S1A-C), but the SEAP activity induced by OMV_47_-LcrV remained substantially higher than that of purified LcrV protein (Fig. S1B). This may be because tetra-acylated diphosphatidylglycerols in outer membranes produced by gram-negative bacteria can activate TLR4/MD-2 depending upon the saturation state of their acyl chains [39]. Another possible reason is that sensing of cytosolic LPS delivered by OMVs via the cysteine protease caspase-11 (caspase-4 and -5 in humans) may activate an NF-κB-inducible SEAP reporter gene in a TLR4-independent fashion [40, 41]. However, the removal of one fatty acid chain from MPLA compromised the immunogenicity of OMV_47_-LcrV when immunized mice were challenged with a high dose of *Y. pestis* (Fig. S1D). Studies have shown that a combination of aluminum salts can alleviate the reactogenicity of vaccines (*Neisseria* OMVs and dTpa) and retain immunogenicity [42, 43]. Thus, we will pursue this strategy to achieve retained immunogenicity but mitigated reactogenicity of OMV_46_-LcrV.

Usually, nonreplicating vaccines administered parenterally fail to elicit mucosal immune responses [44]. However, growing evidence is challenging this concept. Similar to previous studies [45, 46], intramuscular OMV_46_-LcrV immunization resulted in strong LcrV-specific IgG titers in both systemic and lung mucosal compartments (Figs. 3A and S2A). Although no LcrV-specific sIgA was detected in the BALF, OMV_46_-LcrV immunization provided superior protection against pneumonic plague (Figs. 2D and G). Consistent with our previous study [47], the data suggest that sIgA is inessential for protection against pneumonic plague, while IgG matters. To achieve respiratory protection against fully virulent *Y. pestis*, both humoral and cellular immune responses are needed in a synergistic fashion [9, 48]. This perspective was further confirmed by the protective results in mice by antibody transfer and lack of B and/or T cells (Figs. 4B-E).

The hallmark of primary pneumonic plague in naïve mice after 48 h of exposure is functional impairment of AMs along with excessive neutrophil influx into the lungs, which rapidly progresses to death [49]. In addition to the robust induction of adaptive immune responses, OMV_46_-LcrV vaccination heightened the activated AMs and contained excessive neutrophil influx during pulmonary *Y. pestis* infection (Fig. 5). AMs contribute to eliminating pathogens by proinflammatory responses [50–52], as well as suppressing overt neutrophil recruitment and maintaining lung homeostasis by anti-inflammatory responses [53, 54]. Thus, OMV_46_-LcrV immunization appears to reprogram innate cells, including AMs, to develop innate immune memory, i.e., trained immunity [55]. Our previous study showed that T-cell depletion in mice immunized with a live attenuated *Y. pseudotuberculosis* strain reduced the counts of AMs, increased the counts of neutrophils, and impaired protection against pneumonic plague [56]. Here, the lack of B and/or T cells significantly dampened immune protection against pneumonic plague (Figs. 4B-E). Thus, our results imply that the coordination of innate and adaptive immunity in OMV_46_-LcrV-vaccinated animals can effectively halt deadly infection. However, the precise mechanisms by which parenteral OMV_46_-LcrV immunization orchestrates lung immunity remain elusive and need to be investigated further.

In summary, from a vaccine perspective, the new OMV_46_-LcrV exhibited high immunogenicity and low reactogenicity and could be an excellent plague vaccine candidate. Moreover, the YptbS46 strain exclusively producing MPLA can be a platform for delivering foreign antigens against a wide variety of diseases.

## Materials and Methods

### Mice and animal ethics statement

Six-week-old male and female Swiss Webster mice were purchased from Charles River Laboratories (Wilmington, MA) and acclimated for one week before experiments. Wild-type C57/BL6 (B6) and Rag1^−/−^ mice on a B6 background were purchased from Jackson Laboratories (Bar Harbor, Maine, USA). All B6 background mice were bred in an animal facility at Albany Medical College. Age-matched male and female mice were used in the experiments. All animal studies were performed per the NIH “Guide for the Care and Use of Laboratory Animals” and approved by the Institutional Animal Care and Use Committee at Albany Medical College (IACUC protocol# 22-12002).

### Strains, plasmids, culture conditions, molecular procedures, and OMV isolation and analysis

All bacterial strains and plasmids used in this study were listed in Supplementary Information (SI) Table 1. Bacterial cultures, molecular procedures, and OMV isolation and analysis were described in the SI.

### Lipid A isolation and analysis

The detailed procedures for lipid A isolation from Yptb mutants and their OMVs and lipid A analysis using normal-phase liquid chromatography/electrospray ionization-mass spectrometry (NPLC/ESI-MS) were conducted as described previously [10, 57].

### Animal experiments

Groups of mice (7 weeks old, mixed males and females) were primed with OMVs by intramuscular (IM) injection and boosted at week 3 after the initial vaccination. Vaccination with LcrV/Alhydrogel and PBS were used as experimental controls. Blood was collected via the submandibular veins on 21 and 39 DPV to harvest serum for antibody analysis. On 42 DPV, animals were anesthetized with 100 μl of ketamine/xylazine mixture (25 mg/ml ketamine plus 1 mg/ml xylazine) and challenged intranasally with a fully virulent *Y. pestis* KIM6+ (pCD1Ap) strain in 40 μl of PBS to recapitulate pneumonic plague. Animal survival was monitored for 15 days. To determine the bacterial burden after the challenge, animals were euthanized with an overdose of sodium pentobarbital. The lungs, liver, and spleen were collected at 2 DPI and homogenized in ice-cold PBS (pH 7.4) using a bullet blender (Bullet Blender Blue; NY, USA) at power 7 for 2 min. Serial dilutions of each organ homogenate were spread on HIB Congo Red agar plates to enumerate *Y. pestis* colony form units (CFUs).

### Histopathology

Groups of three mice were infected intranasally with 2 × 10^4^ CFU of *Y. pestis* KIM6+(pCD1Ap). At 2 DPI, the lungs were collected from euthanized mice and fixed in 10% formalin overnight before being embedded in paraffin. Lungs from uninfected mice were used as experimental controls. Tissue sections were stained with hematoxylin/eosin and then examined as described previously [26].

### Antibody and cytokine analysis

Anti-LcrV antibody titers in the serum and bronchoalveolar lavage fluid (BALF) were measured using an enzyme-linked immunosorbent assay (ELISA) as described previously [10]. Serum cytokines were measured using commercial mouse cytokine ELISA kits (Invitrogen) according to the manufacturer’s protocol.

### Analysis of T-cell responses in immunized mice before and after infection

Lung and spleen T-cell isolation and activation in immunized mice before and after infection was described previously [56]. To determine *in vitro* activation, single cells (2×10^6^) from the lungs and spleens of euthanized mice were seeded in 24-well cell culture plates and stimulated with endotoxin-free LcrV (20 µg/mL) for 48 h. Then, a 1× brefeldin-A and monensin cocktail (1:1 ratio) was added to block Golgi-mediated cytokine secretion for 2-3 hours before cell collection. To determine *in vivo* activation, single cells (2×10^6^) isolated from the lungs and spleens of euthanized mice at 2 DPI were induced by 50 ng/mL phorbol myristate acetate (PMA) and 1 μg/mL ionomycin for 1 h and then treated with 1× brefeldin-A and monensin cocktail for another 2 h. Induced cells were harvested and resuspended in flow cytometry staining (FACS) buffer containing Fc block (anti-mouse CD16/32 antibodies) (1:200) for 10 min on ice. T cells stained with fluorochrome-labeled anti-CD3, CD4, CD8, IFN-γ, TNF-α, and IL-17A antibodies (SI Table 3) were analyzed using flow cytometry.

### Analysis of alveolar macrophages and neutrophils in the lungs of immunized mice before and after infection

Cells were isolated from the lung and BALF of immunized mice before and after infections as described previously [56] and were stained with Fixable Viability Dye Zombie Red followed by labeling with fluorochrome-labeled anti-CD11b, Ly6G, CD11c, MHCII, F4/80, and Siglec F antibodies (SI Table 3) in FACS buffer. Flow cytometry analysis was performed using BD FACS symphony A3 (BD Biosciences) using FACSDiva software, and data were analyzed using FlowJo v10.7.

### Serum transfer, B-cell depletion, and T-cell depletion

Monoclonal antibodies (mAbs) used to block B or T cells were purchased from Bio X Cell (West Lebanon, NH, USA). B cells were depleted by intraperitoneal (IP) injection with 200 µg of anti-mouse CD20 mAbs (clone MB20-11) 2 days before the initial immunization and booster immunization. To distinguish lung circulating and resident B cells, mice were intravenously (IV) injected via the tail vein with 5 µg of FITC-conjugated anti-CD45.2 mAb diluted in 200 µL sterile PBS. Three minutes post-IV injection, lung single cells were isolated from mice treated with anti-CD20 and isotype control mAbs and stained with fluorochrome-labeled anti-CD19 antibodies for flow cytometry analysis [58]. T cells were depleted by intraperitoneal (IP) injection with 200 µg of anti-mouse CD4 (clone GK1.5) and/or CD8 (Clone 2.430) mAbs at 1 day before *Y. pestis* infection and at 3 DPI. The control mice received an equal amount of isotype control (rat IgG2b mAb Clone-LTF2).

### Statistical analysis

Data were representative of at least two independent experiments. Statistical analyses of comparisons of data among groups were performed with one-way ANOVA/univariate or two-way ANOVA with Tukey post hoc tests. The log-rank (MantelLJCox) test was used for survival analysis. All data were analyzed using GraphPad PRISM 9.0 software. The data are represented as the mean ± standard deviation (ns, no significance; * *P< 0.05*; ** *P< 0.01;* *** *P< 0.001;* **** *P<0.0001*).

## Acknowledgments

The following reagent was obtained through BEI Resources, NIAID, NIH: *Yersinia pestis* LcrV Protein, Recombinant from *Escherichia coli*, NR-32875. This work was supported by the National Institutes of Health grants R01AI162670, R21AI139703, and R01AI125623 to WS, R01AI148366 to ZG, and R01GM143223 to HS. HD thanks the support of NIH supplement for undergraduate summer research experiences under the R01GM143223-02. HS and HD also acknowledge Wadsworth Center’s support of the 3D-EM Facility and assistance from Adam Koplas.

## Contributions

Experiments conceived and designed by: WS and SM; experiments performed by: SM, SD, PL, NY, HD, HS, and ZG; data analyzed by: WS, SM, HS, and ZG; manuscript written by: SM and WS; and manuscript edited by WS, ZG, HS, and HD.

## References

1. Barbieri, R.; Signoli, M.; Cheve, D.; Costedoat, C.; Tzortzis, S.; Aboudharam, G.; Raoult, D.; Drancourt, M., Clin Microbiol Rev 2020, 34 (1). DOI 10.1128/CMR.00044-19.

2. Hinnebusch, B. J., Plague in the 21st Century: Global Public Health Challenges and Goals. Georgiev, V. S., Ed. Humana Press, Totowa, NJ: 2010.

3. Riedel, S., Proc (Bayl Univ Med Cent) 2005, 18 (2), 116–24. DOI 10.1080/08998280.2005.11928049.

4. Rosenzweig, J. A.; Hendrix, E. K.; Chopra, A. K., Appl Microbiol Biotechnol 2021, 105 (12), 4931–4941. DOI 10.1007/s00253-021-11389-6.

5. Perry, R. D.; Fetherston, J. D., Clin Microbiol Rev 1997, 10 (1), 35–66.

6. Galimand, M.; Guiyoule, A.; Gerbaud, G.; Rasoamanana, B.; Chanteau, S.; Carniel, E.; Courvalin, P., N Engl J Med 1997, 337 (10), 677–80.

7. Welch, T. J.; Fricke, W. F.; McDermott, P. F.; White, D. G.; Rosso, M. L.; Rasko, D. A.; Mammel, M. K.; Eppinger, M.; Rosovitz, M. J.; Wagner, D.; Rahalison, L.; Leclerc, J. E.; Hinshaw, J. M.; Lindler, L. E.; Cebula, T. A.; Carniel, E.; Ravel, J., Plos One 2007, 2 (3), e309. DOI 10.1371/journal.pone.0000309 [doi].

8. Kiefer, D.; Dalantai, G.; Damdindorj, T.; Riehm, J. M.; Tomaso, H.; Zoller, L.; Dashdavaa, O.; Pfister, K.; Scholz, H. C., Vector borne and zoonotic diseases 2012, 12 (3), 183–8. DOI 10.1089/vbz.2011.0748.

9. Sun, W.; Singh, A. K., NPJ Vaccines 2019, 4, 11. DOI 10.1038/s41541-019-0105-9.

10. Wang, X.; Li, P.; Singh, A. K.; Zhang, X.; Guan, Z.; Curtiss, R., 3rd; Sun, W., Proc Natl Acad Sci U S A 2022, 119 (11), e2109667119. DOI 10.1073/pnas.2109667119.

11. Majumder, S.; Olson, R. M.; Singh, A.; Wang, X.; Li, P.; Kittana, H.; Anderson, P. E.; Anderson, D. M.; Sun, W., Infect Immun 2022, 90 (8), e0016522. DOI 10.1128/iai.00165-22.

12. Kilgore, P. B.; Sha, J.; Andersson, J. A.; Motin, V. L.; Chopra, A. K., NPJ Vaccines 2021, 6 (1), 21. DOI 10.1038/s41541-020-00275-3.

13. Frey, S. E.; Lottenbach, K.; Graham, I.; Anderson, E.; Bajwa, K.; May, R. C.; Mizel, S. B.; Graff, A.; Belshe, R. B., Vaccine 2017, 35 (48 Pt B), 6759–6765. DOI 10.1016/j.vaccine.2017.09.070.

14. Quenee, L. E.; Ciletti, N. A.; Elli, D.; Hermanas, T. M.; Schneewind, O., Vaccine 2011, 29 (38), 6572–83. DOI 10.1016/j.vaccine.2011.06.119.

15. Aftalion, M.; Tidhar, A.; Vagima, Y.; Gur, D.; Zauberman, A.; Holtzman, T.; Makovitzki, A.; Chitlaru, T.; Mamroud, E.; Levy, Y., Vaccines (Basel) 2023, 11 (3). DOI 10.3390/vaccines11030581.

16. Raetz, C. R.; Whitfield, C., Annu Rev Biochem 2002, 71, 635–700. DOI 10.1146/annurev.biochem.71.110601.135414 [doi] 110601.135414 [pii].

17. Gunn, J. S.; Lim, K. B.; Krueger, J.; Kim, K.; Guo, L.; Hackett, M.; Miller, S. I., Mol Microbiol 1998, 27 (6), 1171–82.

18. Winfield, M. D.; Latifi, T.; Groisman, E. A., J Biol Chem 2005, 280 (15), 14765–72. DOI M413900200 [pii] 10.1074/jbc.M413900200.

19. Knirel, Y. A.; Lindner, B.; Vinogradov, E. V.; Kocharova, N. A.; Senchenkova, S. N.; Shaikhutdinova, R. Z.; Dentovskaya, S. V.; Fursova, N. K.; Bakhteeva, I. V.; Titareva, G. M.; Balakhonov, S. V.; Holst, O.; Gremyakova, T. A.; Pier, G. B.; Anisimov, A. P., Biochemistry 2005, 44 (5), 1731–43. DOI 10.1021/bi048430f [doi].

20. Rebeil, R.; Ernst, R. K.; Gowen, B. B.; Miller, S. I.; Hinnebusch, B. J., Mol Microbiol 2004, 52 (5), 1363–73. DOI 10.1111/j.1365-2958.2004.04059.x [doi] MMI4059 [pii].

21. Sun, W.; Six, D. A.; Reynolds, C. M.; Chung, H. S.; Raetz, C. R.; Curtiss, R., 3rd, Infection and immunity 2013, 81 (4), 1172–85. DOI 10.1128/IAI.01403-12.

22. Johnson, L. E. The pmrHFIJKLM operon in Yersinia pseudotuberculosis enhances resistance to CCL28 and promotes phagocytic engulfment by neutrophils. Brigham Young University, 2016.

23. Raetz, C. R.; Reynolds, C. M.; Trent, M. S.; Bishop, R. E., Annu Rev Biochem 2007, 76, 295–329. DOI 10.1146/annurev.biochem.76.010307.145803 [doi].

24. Bouaziz, J. D.; Yanaba, K.; Venturi, G. M.; Wang, Y.; Tisch, R. M.; Poe, J. C.; Tedder, T. F., Proc Natl Acad Sci U S A 2007, 104 (52), 20878–83. DOI 10.1073/pnas.0709205105.

25. Swanson, R. V.; Gupta, A.; Foreman, T. W.; Lu, L.; Choreno-Parra, J. A.; Mbandi, S. K.; Rosa, B. A.; Akter, S.; Das, S.; Ahmed, M.; Garcia-Hernandez, M. D.; Singh, D. K.; Esaulova, E.; Artyomov, M. N.; Gommerman, J.; Mehra, S.; Zuniga, J.; Mitreva, M.; Scriba, T. J.; Rangel-Moreno, J.; Kaushal, D.; Khader, S. A., Nature Immunology 2023, 24 (5), 855–868. DOI 10.1038/s41590-023-01476-3.

26. Pechous, R. D.; Sivaraman, V.; Price, P. A.; Stasulli, N. M.; Goldman, W. E., PLoS pathogens 2013, 9 (10), e1003679. DOI 10.1371/journal.ppat.1003679.

27. Peters, K. N.; Dhariwala, M. O.; Hughes Hanks, J. M.; Brown, C. R.; Anderson, D. M., PLoS pathogens 2013, 9 (4), e1003324. DOI 10.1371/journal.ppat.1003324.

28. Eichelberger, K. R.; Jones, G. S.; Goldman, W. E., mBio 2019, 10 (6). DOI 10.1128/mBio.02759-19.

29. Kanokudom, S.; Assawakosri, S.; Suntronwong, N.; Chansaenroj, J.; Auphimai, C.; Nilyanimit, P.; Vichaiwattana, P.; Thongmee, T.; Yorsaeng, R.; Duangchinda, T.; Chantima, W.; Pakchotanon, P.; Srimuan, D.; Thatsanatorn, T.; Klinfueng, S.; Mongkolsapaya, J.; Sudhinaraset, N.; Wanlapakorn, N.; Honsawek, S.; Poovorawan, Y., Vaccine 2022, 40 (39), 5657–5663. DOI 10.1016/j.vaccine.2022.08.033.

30. Ellis, T. N.; Kuehn, M. J., Microbiol Mol Biol R 2010, 74 (1), 81–94. DOI 10.1128/Mmbr.00031-09.

31. Santos, J. C.; Dick, M. S.; Lagrange, B.; Degrandi, D.; Pfeffer, K.; Yamamoto, M.; Meunier, E.; Pelczar, P.; Henry, T.; Broz, P., EMBO J 2018, 37 (6). DOI 10.15252/embj.201798089.

32. Acevedo, R.; Fernandez, S.; Zayas, C.; Acosta, A.; Sarmiento, M. E.; Ferro, V. A.; Rosenqvist, E.; Campa, C.; Cardoso, D.; Garcia, L.; Perez, J. L., Front Immunol 2014, 5, 121. DOI 10.3389/fimmu.2014.00121.

33. Kuehn, M. J.; Kesty, N. C., Gene Dev 2005, 19 (22), 2645–2655. DOI 10.1101/gad.1299905.

34. Qing, G.; Gong, N.; Chen, X.; Chen, J.; Zhang, H.; Wang, Y.; Wang, R.; Zhang, S.; Zhang, Z.; Zhao, X.; Luo, Y.; Liang, X., Biophysics Reports 2019, 5, 184–198.

35. van de Waterbeemd, B.; Streefland, M.; van der Ley, P.; Zomer, B.; van Dijken, H.; Martens, D.; Wijffels, R.; van der Pol, L., Vaccine 2010, 28 (30), 4810–6. DOI 10.1016/j.vaccine.2010.04.082.

36. Simpson, B. W.; Trent, M. S., Nat Rev Microbiol 2019, 17 (7), 403–416. DOI 10.1038/s41579-019-0201-x.

37. Needham, B. D.; Carroll, S. M.; Giles, D. K.; Georgiou, G.; Whiteley, M.; Trent, M. S., Proc Natl Acad Sci U S A 2013, 110 (4), 1464–9. DOI 10.1073/pnas.1218080110.

38. Kong, Q.; Six, D. A.; Roland, K. L.; Liu, Q.; Gu, L.; Reynolds, C. M.; Wang, X.; Raetz, C. R.; Curtiss, R. I., J Immunol 2011, 187 (1), 412–23. DOI 10.4049/jimmunol.1100339.

39. Pizzuto, M.; Lonez, C.; Baroja-Mazo, A.; Martinez-Banaclocha, H.; Tourlomousis, P.; Gangloff, M.; Pelegrin, P.; Ruysschaert, J. M.; Gay, N. J.; Bryant, C. E., Cell Mol Life Sci 2019, 76 (18), 3667–3678. DOI 10.1007/s00018-019-03113-5.

40. Vanaja, S. K.; Russo, A. J.; Behl, B.; Banerjee, I.; Yankova, M.; Deshmukh, S. D.; Rathinam, V. A. K., Cell 2016, 165 (5), 1106–1119. DOI 10.1016/j.cell.2016.04.015.

41. Hagar, J. A.; Powell, D. A.; Aachoui, Y.; Ernst, R. K.; Miao, E. A., Science 2013, 341 (6151), 1250–3. DOI 10.1126/science.1240988.

42. Nagaputra, J. C.; Rollier, C. S.; Sadarangani, M.; Hoe, J. C.; Mehta, O. H.; Norheim, G.; Saleem, M.; Chan, H.; Derrick, J. P.; Feavers, I.; Pollard, A. J.; Moxon, E. R., Clin Vaccine Immunol 2014, 21 (2), 234–42. DOI 10.1128/CVI.00561-13.

43. Theeten, H.; Van Damme, P.; Hoppenbrouwers, K.; Vandermeulen, C.; Leback, E.; Sokal, E. M.; Wolter, J.; Schuerman, L., Vaccine 2005, 23 (12), 1515–1521. DOI 10.1016/j.vaccine.2004.08.002.

44. Neutra, M. R.; Kozlowski, P. A., Nat Rev Immunol 2006, 6 (2), 148–58. DOI nri1777 [pii] 10.1038/nri1777.

45. Moldoveanu, Z.; Clements, M. L.; Prince, S. J.; Murphy, B. R.; Mestecky, J., Vaccine 1995, 13 (11), 1006–1012. DOI Doi 10.1016/0264-410x(95)00016-T.

46. Cheng, C. M.; Pal, S.; Bettahi, I.; Oxford, K. L.; Barry, P. A.; de la Maza, L. M., Vaccine 2011, 29 (18), 3456–3464. DOI 10.1016/j.vaccine.2011.02.057.

47. Singh, A. K.; Curtiss, R., 3rd; Sun, W., Infect Immun 2019, 87 (10). DOI 10.1128/IAI.00296-19.

48. Parent, M. A.; Berggren, K. N.; Kummer, L. W.; Wilhelm, L. B.; Szaba, F. M.; Mullarky, I. K.; Smiley, S. T., Infect Immun 2005, 73 (11), 7304–10. DOI 10.1128/IAI.73.11.7304-7310.2005.

49. Pechous, R. D.; Sivaraman, V.; Stasulli, N. M.; Goldman, W. E., Trends Microbiol 2016, 24 (3), 190–197. DOI 10.1016/j.tim.2015.11.008.

50. Janssen, W. J.; Barthel, L.; Muldrow, A.; Oberley-Deegan, R. E.; Kearns, M. T.; Jakubzick, C.; Henson, P. M., Am J Respir Crit Care Med 2011, 184 (5), 547–60. DOI 10.1164/rccm.201011-1891OC.

51. Goritzka, M.; Makris, S.; Kausar, F.; Durant, L. R.; Pereira, C.; Kumagai, Y.; Culley, F. J.; Mack, M.; Akira, S.; Johansson, C., J Exp Med 2015, 212 (5), 699–714. DOI 10.1084/jem.20140825.

52. Aegerter, H.; Kulikauskaite, J.; Crotta, S.; Patel, H.; Kelly, G.; Hessel, E. M.; Mack, M.; Beinke, S.; Wack, A., Nature Immunology 2020, 21 (2), 145–157. DOI 10.1038/s41590-019-0568-x.

53. Zaslona, Z.; Przybranowski, S.; Wilke, C.; van Rooijen, N.; Teitz-Tennenbaum, S.; Osterholzer, J. J.; Wilkinson, J. E.; Moore, B. B.; Peters-Golden, M., J Immunol 2014, 193 (8), 4245–53. DOI 10.4049/jimmunol.1400580.

54. Mould, K. J.; Jackson, N. D.; Henson, P. M.; Seibold, M.; Janssen, W. J., JCI Insight 2019, 4 (5). DOI 10.1172/jci.insight.126556.

55. Netea, M. G.; Joosten, L. A.; Latz, E.; Mills, K. H.; Natoli, G.; Stunnenberg, H. G.; O’Neill, L. A.; Xavier, R. J., Science 2016, 352 (6284), aaf1098. DOI 10.1126/science.aaf1098.

56. Singh, A. K.; Majumder, S.; Wang, X.; Song, R.; Sun, W., J Immunol 2023, 210 (3), 259–270. DOI 10.4049/jimmunol.2200487.

57. Hankins, J. V.; Madsen, J. A.; Needham, B. D.; Brodbelt, J. S.; Trent, M. S., Methods Mol Biol 2013, 966, 239–258. DOI 10.1007/978-1-62703-245-2_15.

58. Barker, K. A.; Etesami, N. S.; Shenoy, A. T.; Arafa, E. I.; Lyon de Ana, C.; Smith, N. M.; Martin, I. M.; Goltry, W. N.; Barron, A. M.; Browning, J. L.; Kathuria, H.; Belkina, A. C.; Guillon, A.; Zhong, X.; Crossland, N. A.; Jones, M. R.; Quinton, L. J.; Mizgerd, J. P., J Clin Invest 2021, 131 (11). DOI 10.1172/JCI141810.

