## Supplementary File for "Pneumonic plague protection induced by a monophosphoryl lipid A decorated *Yersinia* outer-membrane-vesicle vaccine"

Running Title:

***Yersinia* outer membrane vesicle (OMV) vaccine against pneumonic plague**

**Conflict of interest:** All other authors declare that they have no conflicts of interest.

#### MATERIALS AND METHODS

**Bacterial culture conditions.** *E. coli* strains were grown routinely at 37°C in LB broth or LB agar (Difco) [1]. *Y. pseudotuberculosis* (Yptb) strains were grown at 28°C in LB broth and LB agar [2]. The *E. coli* strain χ7213 [3] was used to construct suicide vectors and conjugated with a Yptb strain to generate corresponding mutations. Diaminopimelic acid (DAP) at 50 µg/ml, ampicillin at 100 µg/ml, or chloramphenicol at 25 µg/ml was added to the medium when necessary.

Single colonies of *Y. pestis* KIM6+ (pCD1Ap) grown on heart infusion broth (HIB) Congo red agar plates were used to inoculate HIB supplemented with ampicillin (final concentration 100 µg/ml) and grown overnight at 26°C. Bacteria were then inoculated into 10 ml of fresh HIB enriched with 0.2% xylose and 2.5 mM $CaCl_2$ to obtain an $OD_{620}$ of 0.1 and incubated at 37°C to reach an $OD_{620}$ of 0.6. Bacteria were pelleted and resuspended in 1 ml of isotonic PBS and then adjusted to an appropriate concentration for the intranasal challenge [4]. Bacterial titers were determined by serial dilution, plating, and counting colonies.

**Molecular and genetic procedures.** The plasmids and primers used in this study are listed in Supplementary Information (SI) Tables 1 and 2, respectively. To generate the *pmrF-J* mutation, the Δ*pmrF-J* flanking region of Yptb PB1+ was assembled by overlapping PCR using PmrF-J-1/PmrF-J-2 and PmrF-J-3/PmrF-J-4 primer sets and cloned into the suicide vector pRE112 to generate pSMV65 (Table 1). The suicide plasmid pSMV66 for the *lpxL* deletion was constructed following the same procedures. All plasmids were confirmed by PCR screening and DNA sequencing. The procedures for the *sacB*-based sucrose counterselectable suicide vectors used to construct unmarked deletion mutations in Yptb were described in our previous report [2]. Successful gene mutations were confirmed by PCR screening and DNA sequencing.

**OMV isolation and analysis.** OMVs were isolated from Yptb mutant strains harboring an Asd+ plasmid as described previously [4]. Briefly, strains were grown at 28°C in LB broth for 24 hr. The bacterial cultures were kept on ice for 2 hr. Then, the bacterial cells were pelleted by centrifugation at 10,000 × g and 4°C for 20 min. The culture supernatant was filtered using a 0.45-μm pore membrane (Millipore) to remove any residual bacteria and cell debris and concentrated with a 100-kDa filter using a Vivaflow 200 system (Sartorius). OMVs were harvested by ultracentrifugation (120,000 × g) for 2 h at 4°C. The vesicle pellet was washed and resuspended in 0.1× sterilized PBS (pH 7.4), and the ultracentrifugation step was repeated. The final vesicle pellet was resuspended in 0.1× sterilized PBS, filtered through a 0.22-μm pore membrane (Millipore), and stored at 4°C for subsequent experiments.

The total protein and lipid contents of OMVs were routinely measured using a Bradford assay and an FM4-64 fluorescent dye-binding assay as described previously [4]. The major antigens present in OMV preparations were detected by western blotting. The size and morphology of the OMVs were characterized using DynaPro™ Dynamic Light Scattering (DLS) (Wyatt Technology, Santa Barbara, CA, USA) and a JEOL 1200EX transmission electron microscope (JEOL Peabody, MA) equipped with an AMT 8-megapixel digital camera (Advanced Microscopy Techniques, Woburn, MA). OMVs were analyzed by TEM as described in previous reports with modificaiotn[5]. In brief, 3 uL of the samples were applied to glow-discharged carbon-film-coated EM grids for 1 minute before a staining by 3 uL 3% uranyl acetate for 45 seconds. After revmoved extra staining solution, the grids were air-dried for at least one hour before examed uisng a Tecnai F20 electron microscope (FEI, Hillsboro, OR, USA) operated at 200 kV. Micrographs were recorded using a 4K x 4K CMOS CCD camera (TVIPS

Temcam F416). Furthermore, the TLR4 stimulatory activity of OMVs was assayed using HEK-Blue$^{TM}$ mTLR4 cells (Invivogen, CA, USA) as described previously [6].

**Table 1. Strains and plasmids used in this study**

| Strain or Plasmid | Genotype or relevant characteristics | Source |
|---|---|---|
| Strains | | |
| *E. coli* χ6212 | *F– λ– ϕ80 Δ(lacZYA-argF) endA1 recA1 hsdR17 deoR thi-1 glnV44 gyrA96 relA1 ΔasdA4* | [7] |
| *E. coli* χ7213 | *thi-1 thr-1 leuB6 fhuA21 lacY1 glnV44 ΔasdA4 recA1* RP4 2-Tc::Mu [λ*pir*]; $Km^r$ | [7] |
| ***Y. pestis*** | | |
| KIM6+ (pCD1Ap) | pCD1Ap, pMT1, pPCP1, $Pgm^+$, wild-type strain | [8] |
| *Y. pseudotuberculosis* PB1+ | Serotype O:1B | [9] |
| YptbS44 | Δ*tolR* Δ*asd* Δ*lacI* :: $P_{lpp}$ *lpxE* Δ*hmsHFRS* $pYV^-$ Δ*lacZ* :: *caf1R-caf1M-caf1A-caf1* | [6] |
| YptbS46 | Δ*pmrF-J* Δ*tolR* Δ*asd* Δ*lacI* :: $P_{lpp}$ *lpxE* Δ*hmsHFRS* $pYV^-$ Δ*lacZ* :: *caf1R-caf1M-caf1A-caf1* | This study |
| YptbS47 | Δ*lpxL* Δ*pmrF-J* Δ*tolR* Δ*asd* Δ*lacI* :: $P_{lpp}$ *lpxE* Δ*hmsHFRS* $pYV^-$ Δ*lacZ* :: *caf1R-caf1M-caf1A-caf1* | This study |
| **Plasmids** | | |
| pRE112 | Suicide vector, $Cm^r$, *mob*$^-$ (RP4)R6K *ori*, *sacB* | [10] |
| pYA3493 | $Asd^+$; β-lactamase signal sequence-based periplasmic secretion, pBR *ori* | [4] |
| pSMV13 | The full-length *Y. pestis lcrV* was cloned into pYA3620 | [4] |

| | | |
|---|---|---|
| pSMV65 | The flanking regions of Δ*pmrF-J* of *Y. pseudotuberculosis* into *Xma*I and *Kpn*I sites of pRE112 | This study |
| pSMV66 | The flanking regions of Δ*lpxL* of *Y. pseudotuberculosis* into *Xma*I and *Kpn*I sites of pRE112 | This study |

¶: $Cm^{r}$, chloramphenicol resistance; Ap, ampicillin resistance.

**SI Table 2. Primers used in this work**

| Name | Sequence |
|---|---|
| PmrF-J-1 | 5’cggggtaccatgcagagttttttgcctttttcta3’ |
| PmrF-J-2 | 5’ gctgttttaccgcggcgcgccagcatagtggacgggaa3’ |
| PmrF-J-3 | 5’ atgctggcgcgccgcggtaaaacagcaattcgatttta3’ |
| PmrF-J-4 | 5’cggcccgggtcattcagcatcctttatgtctga3’ |
| LpxL-1 | 5’cgggagctcgcgcaccggagtcttcaccacctt3’ |
| LpxL-2 | 5’cgcgtccggaatgcccgtccggacggttccgacgacgatca3’ |
| LpxL-3 | 5’tgatcgtcgtcggaaccgtccggacgggcattccggacgcg3’ |
| LpxL-4 | 5’cggtctagatcgccgaagtactcgcggttga3’ |

**SI Table 3. Flow cytometry antibodies used in this study**

| Name | Vendor | Clone No. |
|---|---|---|
| Live Dead- Zombie Red | BioLegend | |
| CD3- PECy7 or allophycocyanin-Cy7 | BD | 145-2C11 |
| CD4-PE | BioLegend | B359231 |
| IFN-γ- APC | BioLegend | XMG1.2 |
| TNF-α- BV510 | BioLegend | MP6-XT22 |
| IL-17A-APC-Cy7/Allophycocyanin | BioLegend | TC11-18H10.1 |
| CD11b- BV510 | BioLegend | M1/70 |
| Ly6G-PECy7 | BioLegend | 1A8 |
| CD11C-APC-Cy7 | BioLegend | N418 |
| MHCII-BV421 | BioLegend | M5/114.15.2 |
| F4/80-APC | BioLegend | BM8 |
| SiglecF-A700 | Invitrogen | 1RMN44N |
| CD45.2-FITC | BioLegend | S18009F |
| CD19b-APC-cy7 | BioLegend | 6D5 |

**SI Figure legends**

**Fig. S1. Lipid profiles of the Yptb47 strain, TLR4 stimulatory activity of $OMV_{47}$-LcrV, and protective evaluation of $OMV_{47}$-LcrV immunization.** (A) Mass spectrometry analysis of lipid A species in the Yptb47 strain (Table S1). Shown are the negative ion electropray ionization (ESI) mass spectrum of the $[M\text{-}H]^-$ ions of tri-acylated and tera-acylated lipid A species and their chemical structures. (B) Comparison of the secreted embryonic alkaline phosphatase (SEAP) activities in HEK-Blue™ cells with or without murine TRL4. HEK-Blue™ mTLR4 (InvivoGen) cells were cocultured with 40 µg/ml $OMV_{44}$-LcrV, $OMV_{46}$-LcrV, $OMV_{47}$-LcrV, or purified LcrV for 8 h. (C) Weight change rates of mice after immunization with 30 µg of $OMV_{46}$-LcrV/50 µl of PBS, 30 µg of $OMV_{47}$-LcrV/50 µl of PBS, and 50 µl of PBS (negative control). (D) On day 42 after the initial immunization, $OMV_{47}$-LcrV-immunized Swiss Webster mice were intranasally challenged with 200 and 2,000 $LD_{50}$ of *Y. pestis* KIM6+(pCD1Ap). Morbidity and mortality were recorded for 15 days. Data are shown as the mean ± SD. The statistical significance of differences among groups was analyzed by two-way multivariate ANOVA with Tukey's post hoc test. Statistical significance was analyzed by the log-rank (Mantel-Cox) test for survival analysis. ns, no significance; *, $P< 0.05$; **, $P< 0.01$; ***, $P< 0.001$; ****, $P< 0.0001$.

**Fig. S2. Antibody titer in BALF, T-cell gating, and spleen antigen-specific T-cell responses.** (A) Anti-LcrV IgG titers in the BALFs of $OMV_{46}$-LcrV-, $OMV_{46}$-NA-, rLcrV-, or PBS-immunized mice. On 41 DPV, immunized mice were euthanized, and the BALFs were collected immediately. (B) Gating Strategy for T-cell responses. (C) LcrV-specific spleen T-cell responses in immunized mice before *Y. pestis* infection. (D) Spleen T-cell responses in immunized mice at 2 DPI. The detailed procedure is described in the Materials and Methods. Each symbol was obtained from an individual mouse, and data were represented as the mean ± SD. The statistical significance of differences among groups was analyzed by two-way ANOVA with Tukey's post hoc test: ns, no significance; *, *P< 0.05*; **, *P< 0.01*; ****, *P< 0.0001*.

**Fig. S3. T-cell depletion, B-cell depletion, and gating strategy for lung myeloid cells.** (A) Flow plots for $CD4^+$ and $CD8^+$ T-cell depletion. (B) Gating strategy for B-cell depletion. (C) Flow and quantitative plots for B-cell depletion. Swiss Webster mice (n=5/group, females) were intravenously (IV) injected via the tail vein with 5 µg of FITC-conjugated anti-CD45.2 mAb to distinguish lung circulating and resident B cells as described in the Materials and Methods. (Left) Flow plots showing circulating and resident B cells in the lungs of mice treated with anti-CD20 or isotype control mAbs; (right) a quantitative plot for circulating and resident B cells. (D) Serum anti-LcrV titers in $OMV_{46}$-LcrV-immunized mice treated with anti-CD20 or isotype control mAbs. (E) Gating strategy for myeloid cells in the lung. Each symbol was obtained from an individual mouse, and data were represented as the mean ± SD. The statistical significance of differences among groups was analyzed by two-way ANOVA with Tukey's post hoc test: ns, no significance; *, *P< 0.05*; **, *P< 0.01*; ****, *P< 0.0001*.

#### References

1. Bertani, G., *J Bacteriol* **1951,** *62* (3), 293-300.

2. Sun, W.; Sanapala, S.; Henderson, J. C.; Sam, S.; Olinzock, J.; Trent, M. S.; Curtiss, R., 3rd, *Infection and immunity* **2014,** *82* (10), 4390-404. DOI 10.1128/IAI.02173-14.

3. Roland, K.; Curtiss, R. I.; Sizemore, D., *Avian diseases* **1999,** *43* (3), 429-41.

4. Wang, X.; Singh, A. K.; Zhang, X.; Sun, W., *Infect Immun* **2020,** *88* (5), e00081-20. DOI 10.1128/IAI.00081-20.

5. Eddy, J. L.; Gielda, L. M.; Caulfield, A. J.; Rangel, S. M.; Lathem, W. W., *Plos One* **2014,** *9* (9). DOI ARTN e107002 10.1371/journal.pone.0107002.

6. Wang, X.; Li, P.; Singh, A. K.; Zhang, X.; Guan, Z.; Curtiss, R., 3rd; Sun, W., *Proc Natl Acad Sci U S A* **2022,** *119* (11), e2109667119. DOI 10.1073/pnas.2109667119.

7. Kang, H. Y.; Srinivasan, J.; Curtiss, R., III, *Infection and immunity* **2002,** *70* (4), 1739-49.

8. Sun, W., Six, D., Kuang, X. Y., Roland, K. L., Raetz, C. R. H., Curtiss, R. III., *Vaccine* **2011,** *29*, 2986-2998.

9. Sun, W.; Sanapala, S.; Rahav, H.; Curtiss, R., 3rd, *Vaccine* **2015,** *33*, 6727-35. DOI 10.1016/j.vaccine.2015.10.074.

10. Edwards, R. A.; Keller, L. H.; Schifferli, D. M., *Gene* **1998,** *207* (2), 149-157. DOI Doi 10.1016/S0378-1119(97)00619-7.

Figures:

Figure S1:

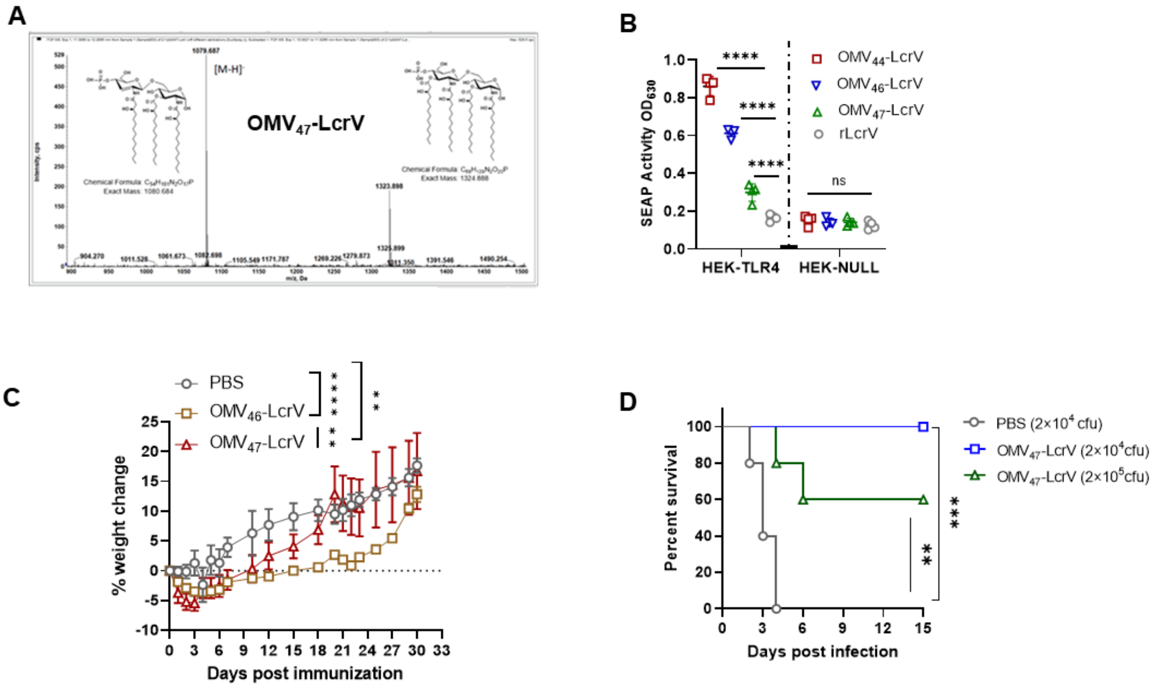


**Supplementary Figure 1**

Figure S2:

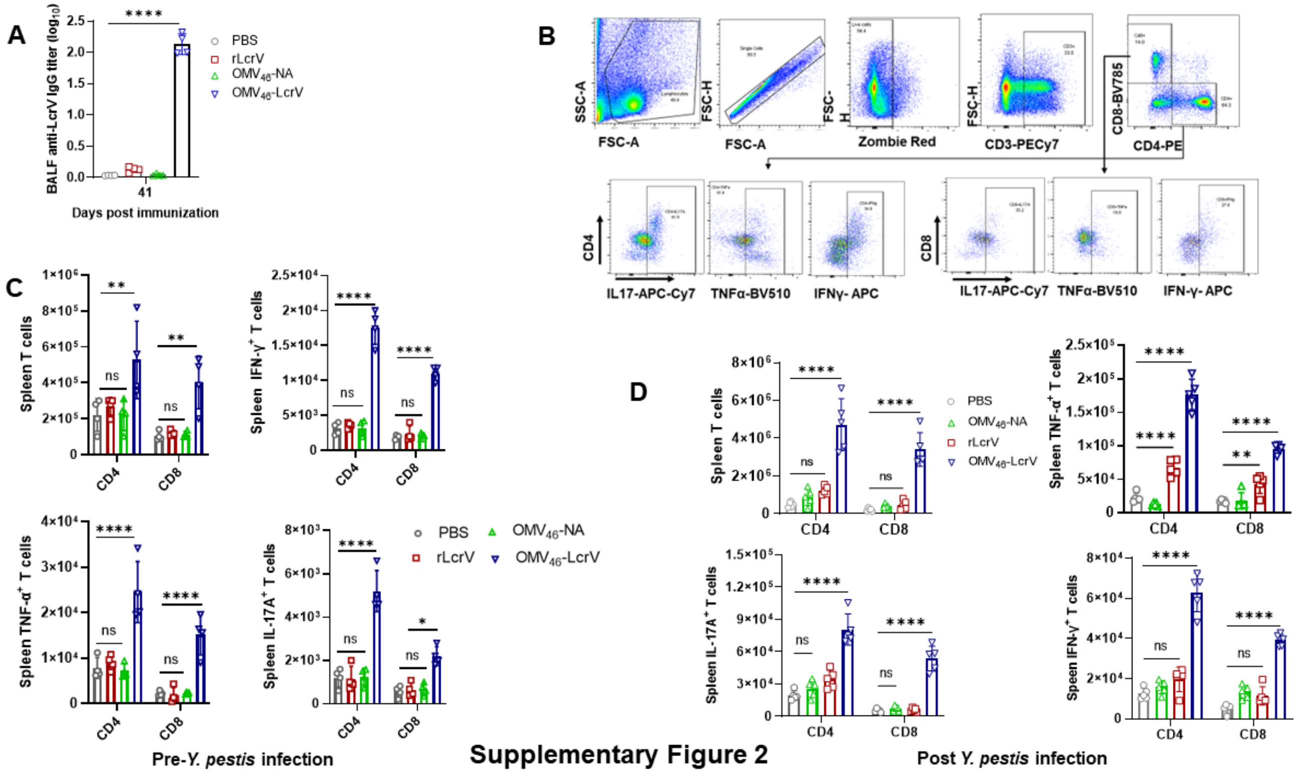


**Supplementary Figure 2**

Figure S3:

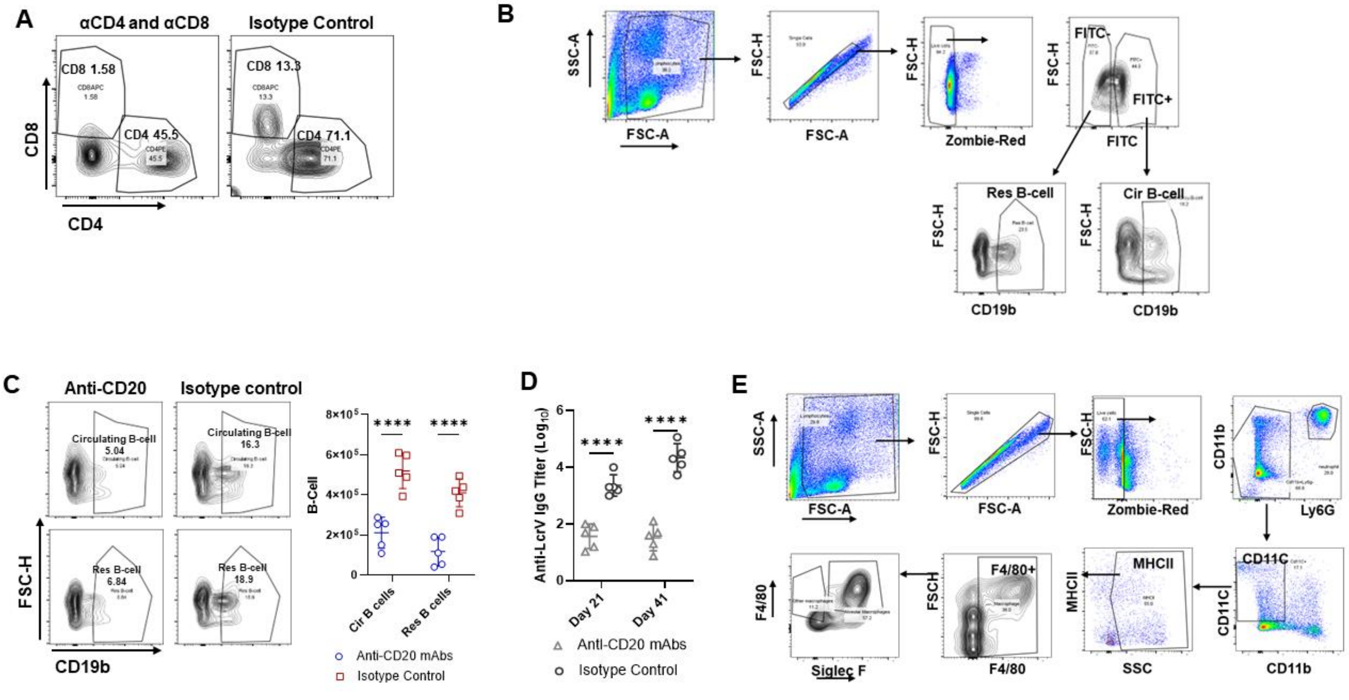


**Supplementary Figure 3**
